# The genome sequence of *Sorbus pohuashanensis* provides insights into population evolution and leaf sunburn response

**DOI:** 10.1101/2021.08.27.457897

**Authors:** Dongxue Zhao, Yan Zhang, Yizeng Lu, Liqiang Fan, Zhibin Zhang, Mao Chai, Jian Zheng

**Affiliations:** School of Landscape Architecture, Beijing University of Agriculture, Beijing, 102206, China; Shandong Provincial Center of Forest Tree Germplasm Resources, Jinan, Shandong Province, 250102, China; Institute of Cotton Research of the Chinese Academy of Agricultural Sciences, Anyang, Henan province, 455000, China

**Keywords:** *Sorbus pohuashanensis*, genome assembly, population evolution, sunburn

## Abstract

*Sorbus pohuashanensis* is a potential horticulture and medicinal plant, but its genomic and genetic background remains unknown. Here, we de novo sequenced and assembled the *S. pohuashanensis* (Hance) Hedl. reference genome using PacBio long reads. Based on the new reference genome, we resequenced a core collection of 22 *Sorbus* spp. samples, which were divided into two groups (G1 and G2) based on phylogenetic and PCA analysis. These phylogenetic clusters were highly in accordance with the classification based on leaf shape. Natural hybridization between the G1 and G2 groups was evidenced by a sample (R21) with a highly heterozygous genotype. Nucleotide diversity (π) analysis showed that G1 has a higher diversity than G2, and that G2 originated from G1. During the evolution process, the gene families involved in photosynthesis pathways expanded and gene families involved in energy consumption contracted. Comparative genome analysis showed that *S. pohuashanensis* has a high level of chromosomal synteny with *Malus domestica* and *Pyrus communis*. RNA-seq data suggested that flavonol biosynthesis and heat-shock protein (HSP)-heat-shock factor (HSF) pathways play important roles in protection against sunburn. This research provides new insight into the evolution of *Sorbus* spp. genomes. In addition, the genomic resources and the identified genetic variations, especially those genes related to stress resistance, will help future efforts to introduce and breed *Sorbus* spp.

## Introduction

*Sorbus* is a small genus of Rosaceae (subfamily Maloideae) composed of more than 250 species that are widely distributed in Asia, Europe, and North America. *Sorbus pohuashanensis* (Hance) Hedl. is widely distributed in North China, Northwest China, and Northeast China. It prefers to grow on slopes or valley forests at an altitude of 900–2500 m and is resistant to low temperatures (Ze Zhang et al., 2020). *S. pohuashanensis* is the tree that can attract birdlife, thus increasing biodiversity and providing considerable ecological benefits (Fig. 1).

**Fig. 1.**
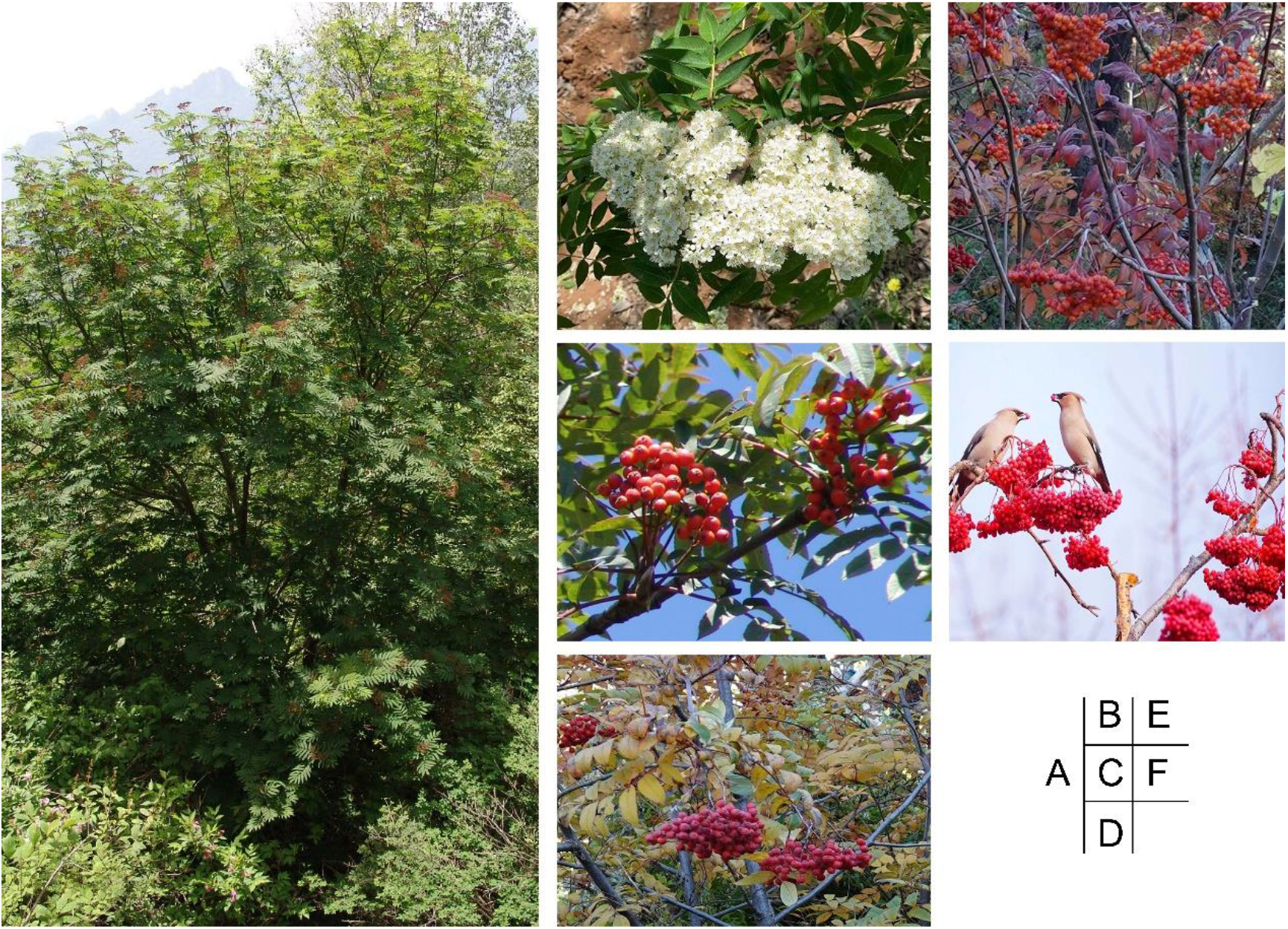
Images of *Sorbus pohuashanensis*. *S. pohuashanensis* mature tree in summer. **B**. flowers in spring. **C**. fruits in early autumn. D, E. fruits in middle autumn. **F**. fruits in late autumn.

It has been recorded in the Chinese Materia Medica that *Sorbus* fruits relieve cough and resolve phlegm, and strengthen the spleen. In recent years, medicinal active ingredients including flavonoids, triterpenoids, anthocyanins, cyanogenic glycosides, and quinones have been isolated from *S. pohuashanensis* and demonstrated to have potential health and medicinal value (Yin et al., 2019; X. Yu et al., 2017).

Because the color of its leaves, fruits and flowers changes with the seasons, *S. pohuashanensis* has attracted the attention of gardeners as an ornamental plant (Fig. 1). Scientists have tried to introduce *S. pohuashanensis* to the plains of China as a horticultural crop. However, in the summer when plants are exposed to sunshine, necrosis spots are generated on the leaves, a phenomenon that has been called sunburn (Munné-Bosch & Vincent, 2019). Sunburn is one of the most serious problems facing the application of *S. pohuashanensis* and other plants. Sunburn can occur in many tissues or organs such as fruits, leaves, and branches, and has been reported in *P. communis* (Goodwin, McClymont, Turpin, & Darbyshire, 2018), *Eriobotrya japonica* and *M. domestica* (Racsko & Schrader, 2012), *Vitis vinifera* (Rustioni et al., 2020), and *Citrus reticulata* Blanco (El-Tanany, Kheder, & Abdallah, 2019). Sunburn not only affects crop yield but also reduces the ornamental value, resulting in millions of dollars in economic losses to growers annually. It has been shown that the occurrence of sunburn is related to many meteorological factors such as light intensity, temperature, air humidity, and wind speed (Racsko & Schrader, 2012). However, little is known about the causes of sunburn in *S. pohuashanensis*.

Long-read sequencing technology has been widely used in many crops such as rice (Du et al., 2017), maize (Jiao et al., 2017) and cotton (Zhaoen Yang et al., 2019), and has promoted basic and applied plant research. The genome sequences of domesticated apple (*M. domestica*) (Sun et al., 2020) and pear (*P. communis*), which belong to the subfamily Maloideae, have also been reported. Here, we report the first reference genome for *S. pohuashanensis*, obtained using PacBio circular consensus sequencing and high throughput chromosome conformation capture (Hi-C) technology. We resequenced a core collection of *Sorbus* spp., mainly from China, and analyzed their phylogenetic relationships. We found a natural hybridization between simple-leaf species and compound-leaf species. We also performed gene family expansion, comparative genomic, and RNA-seq analyses to elucidate the possible mechanisms of sunburn phenomenon.

## Materials and methods

### Sampling for resequencing

We collected 22 *Sorbus* L. samples from 11 species that mainly from China. Including seven Sorbus compound-leaf species, *Sorbus pohuashanensi* (R05-R14, R21, and R22), *Sorbus discolor* Maxim. (R01), *Sorbus aucuparia* Linn. (R17), *Sorbus americana* (R19), *Sorbus hupehensis* Schneid. (R15), *Sorbus discolor* var. *paucijuga* D.K.Zang et P.C.Huang (R16) and *Sorbus aucuparia* ‘Titan’ (R02). The other four species were *Sorbus* simple-leaf, *Sorbus alnifolia* (Sieb. et Zucc.) K. Koch (R03), *Sorbus caloneura* Rehd. (R04), *Sorbus hemsleyi* (R18), and *Sorbus keissleri* (C. K. Schneid.) Rehder (R20) for resequencing (Supplementary Table 5).

### Sampling for transcriptome analysis

The *S. pohuashanensis* plants used in this experiment were compound-leaf types from Beijing and planted in the forestry resource nursery of the School of Landscape and Architecture, Beijing University of Agriculture. A total of 18 well-grown, stable, pest-free 2-year-old seedlings were selected and planted in plastic pots of 24 cm in diameter and 20 cm in depth. All the seedlings were divided into three groups with six plants in each group and placed under a shade awning with 70% shading in March 2020, until the first group sample was taken in the shade awning as control at 14:30 on June 14, 2020. After the first group of samples were taken, they were immediately put into the field. This moment was used as the starting time (0h). The samples have been kept in the field since then, and the sample taken was also completed in the field. After 2.5h, 26.5h, 146.5h from the start time, three group samples were taken as three treatments, which corresponds to level1, level2, and level3. Three biological replicates were set for each group samples. The corresponding temperature, light intensity, and humidity for each group sampling were shown in Supplementary Table 6. For each sample, one leaflet of the fifth compound leaf counting from the top of the plant was collected. The leaves of each group were put immediately in liquid nitrogen and kept in a freezer at -80°C.

### PacBio library construction and sequencing

All procedures were conducted according to the standard protocol supplied by PacBio. Library construction consisted of the following steps: shearing the DNA sample using g-TUBE; repairing the damaged DNA; repairing the DNA termini; connecting dumbbell-shaped adapters; performing exonuclease digestion, and target fragment screening using BluePippin to obtain a sequencing library.

### Hi-C library construction and sequencing

We constructed Hi-C fragment libraries as reported by Fu et al.(Fu et al., 2021). The parameters of LACHESIS were set as follows: C-LUSTER_MIN_RE_SITES, 68; CLUSTER_MAX_LINK_DENSITY, 2; ORDER_MIN_N_RES_IN_TRUN, 55; ORDER_MIN_N_RES_IN_SHREDS, 58. The clean Hi-C reads, accounting for 100-fold coverage of the *S. pohuashanensis* genome. Finally, 404 scaffolds, accounting for 97.29% of the total length, were anchored to the chromosomes.

### Repeat sequence annotation

LTR_FINDER (Z. Xu & Wang, 2007) and RepeatScout (Price, Jones, & Pevzner, 2005) software were employed to construct the database of repeat sequences of the genome. PASTEClassifier (Hoede et al., 2014) was used to classify the database. The resulting database was merged with the Repbase database (Jurka et al., 2005) and used as the final database of repeat sequences, and then based on this database to predict the repeat sequences in the genome using RepeatMasker (Tarailo - Graovac & Chen, 2009) software.

The parameters: LTR_FINDER, RepeatScout, and PASTEClassifier used the default parameters. The RepeatMasker parameters were: -nolow -no_is -norna -engine wublast.

### Gene prediction and function annotation

De novo prediction, homology-based prediction, and UniGene prediction were three approaches to predict coding genes as described by Fu et al.(Fu et al., 2021). For the de novo prediction with the software, default parameters were used. The Hisat parameters were as follows: --max-intronlen 20000, --min-intronlen 20. For Stringtie, TransDecoder, GeneMarks-T, and EVM the default parameters were used. For PASA the parameters were as follows: -align_tools gmap, -maxIntronLen 20000.

The predicted gene sequences were used as queries in BLAST (Altschul, Gish, Miller, Myers, & Lipman, 1990) v2.2.31 (e-value 1e-5) search against the NR (Marchler-Bauer et al., 2010), KOG (Koonin et al., 2004), GO (Dimmer et al., 2012), KEGG (Kanehisa & Goto, 2000), and TrEMBL (Boeckmann et al., 2003) databases. Gene function annotation analysis was conducted with gene KEGG pathway annotations, KOG function annotations, and GO function annotations.

### Whole-genome duplication analysis

We used the stricter version of DupGen_finder (Qiao et al., 2019) GenDup_finder-unique with default parameters to identify genes derived from different modes of gene duplication: WGD, TD, PD (separated by fewer than ten genes on the same chromosome), TRD, and DSD. clusterProfiler v3.14.0 (G. Yu, Wang, Han, & He, 2012) was used for GO and KEGG enrichment analysis of gene families expanded via different duplication modes.

ParaAT 2.0 (Zhang Zhang et al., 2012) was used to calculate the *Ka, Ks*, and *Ka/Ks* values of gene pairs derived from different duplication modes by calling KaKs_Calculator v2.0 software (D. Wang, Zhang, Zhang, Zhu, & Yu, 2010).

WGD v1.1.1 software was used to identify five species of Rosaceae (*F. vesca, P. persica, P. communis, M. domestica and S. pohuashanensis*) ancient whole-genome duplications. Simultaneously use WGD to calculate Ks for five species (Zwaenepoel & Van de Peer, 2018). Perl script was used to calculate the proportion of each homologous gene to the 4DTv site.

### Analysis of LTR insertion time

For analysis of LTR transposon insertion times, LTR_finder v1.07 software (Z. Xu & Wang, 2007) was used to find LTR sequences with scores greater than or equal to 6 points in the genome (parameter: -S 6). The LTR sequences were extracted and compared with MAFFT (parameters: --localpair -- maxiterate 1000), and the Kimura model in EMBOSS v6.6.0 (Rice, Longden, & Bleasby, 2000) was to calculate the distance (K). The formula for calculating the time was T = K/(2 × r), where the molecular clock r value selected was 7*10^−9^ (Ossowski et al., 2010).

LTR_finder V1.07 software (parameters: -D 15000 -L 7000 -C -M 0.9) and LTRharvest V1.6.1 software (Ellinghaus, Kurtz, & Willhoeft, 2008) (parameters: -similar 90 -vic 10 -seed 20 -seqids yes - minlenltr 100 -maxlenltr 7000 -mintsd 4 -maxtsd 6 -motif TGCA -motifmis 1) were performed to identify full-length LTR repeat retrotransposons (LTR-RTs) in the genome. LTR_retriever V2.9.0 software (Ou & Jiang, 2018) with default parameters was used to combine the LTR retrotransposons identified by LTR_finder and LTRhavest, and remove duplicates, determine the only LTR transposon, and calculate the LAI value. The distribution of LTR transposons on the genome windowsize was set to 3Mb. Each point represents the LAI score for a sliding window of 3 Mb in 300-Kb increment.

### Phylogenetic analysis and estimation of species divergence time

IQ-TREE (v1.6.11) (Nguyen, Schmidt, von Haeseler, & Minh, 2014) was applied to construct a phylogenetic tree with 1241 (defined: 80.0% of species having single-copy) single-copy gene sequences. IQ-TREE’s built-in model detection tool ModelFinder (Kalyaanamoorthy, Minh, Wong, von Haeseler, & Jermiin, 2017) obtain the best model was JTT+F+I+G4. This best model was used to construct an evolutionary tree by the maximum likelihood (ML) method, and the bootstraps was set to 1000.

The package MCMCTREE in PAML v4.9i software (Ziheng Yang, 1997) was used to calculate the divergence time, and *A. trichopoda* was used as the outgroup of the root tree. Using TimeTree (http://www.timetree.org/), the divergence times were estimated as follows: *A. trichopoda* vs *P. trichocarpa* at 168–194 MYA, *V. vinifera* vs *G. hirsutum* at 105–115 MYA, *S. pohuashanensis* vs *M domestica* at 10.9–43 MYA. The module MCMCTREE in PAML was then used to estimate the gradient and Hessian parameters required for calculating the divergence time. Finally, the maximum likelihood method and correlated molecular clock and JC69 model were used to estimate the divergence time. Two repeated calculations were performed to determine the consistency between estimates. MCMCTreeR (v1.1) (Puttick, 2019) was used to visualize the evolutionary trees with differentiation time graphically.

### Gene family classification and enrichment analysis

The protein sequences of 10 species (*V. vinifera, F. vesca, P. persica, P. trichocarpa, G. hirsutum, A. thaliana, P. communis, M. domestica, A. trichopoda*, and *S. pohuashanensis*) were used to carry out family classification by Orthofinder v2.4 software (Emms & Kelly, 2019) (diamond comparison method, e value 0.001). The obtained gene families were annotated using the PANTHER V15 database (Mi, Muruganujan, Ebert, Huang, & Thomas, 2018). ClusterProfiler v3.14.0 (G. Yu et al., 2012) was employed to conduct GO and KEGG enrichment analysis of gene families specific to *S. pohuashanensis*.

### The expansion and contraction of gene family

By using CAFE v4.2 software (Han, Thomas, Lugo-Martinez, & Hahn, 2013), the results of phylogenetic tree with divergence time and gene family clustering were used to predict the shrinkage and expansion of the species’ gene families relative to their ancestors. The criteria for defining significant expansion or contraction were a family-wide P-value < 0.05 and a viterbi P-value < 0.05.

### Positive selection analysis

The positive selection analysis was performed using the CodeML module in PAML. Specifically, the single-copy gene families in *F. vesca, P. communis, M. domestica, S. pohuashanensis*, and *P. persica* were obtained. Then each gene family was analyzed using MAFFT (parameters: --localpair --maxiterate 1000). The protein sequence was aligned, and then PAL2NAL was used to invert the codon alignment sequence. Finally, CodeML (using the F3×4 model of codon frequencies) was used to obtain positively selected genes.

### Collinearity analysis

Gene sequences of two species were compared by Diamond (v0.9.29.130) (Buchfink, Xie, & Huson, 2015) (parameter: e<1e−5) to identify similar gene pairs. The C score value was used to filter the blast results by JCVI v0.9.13 (Tang, Krishnakumar, Li, & Zhang, 2015) (parameter: C-score>0.5). MCScanX (Y. Wang et al., 2012) (parameter: -m 5) was used to obtain all the genes in collinear blocks from a gff3 file. JCVI was used to draw the collinearity picture of the linear pattern of each species. VGSC (Y. Xu et al., 2016) was used to show the results of collinearity analysis in the form of dot and bar graphs.

### Illumina short-read sequencing

Genomic DNA was extracted from a single plant of each accession using a Plant DNA Mini Kit (Aidlab Biotech). DNA libraries with 350-bp inserts were constructed for each accession using the Illumina NovaSeq 6000 platform following the manufacturer’s specifications, and 125-bp paired-end reads were generated.

To improve the reliability of the clean data, we excluded the following types of reads described in Ma et al. (Ma et al., 2018). Consequently, 178.71 Gb high-quality clean data were used for subsequent analysis.

### Read alignment, variation calling, and annotation

The remaining clean paired-end reads were aligned to the reference genome of *S. pohuashanensis* with BWA (v0.7.17) with the command ‘mem -t 10 -k 32 -M’. BAM alignment files were then generated by SAMtools (Li et al., 2009) (v1.9, settings: -bS –t). SNP and Indel calling and filtering were conducted with the GATK v4.1.3.0 (DePristo et al., 2011) best practice. Use the ANNOVAR tool with default parameters to annotate SNPs and Indels further (H. Yang & Wang, 2015).

### Phylogenetic tree and population structure

An individual-based NJ tree was constructed based on *P* distance using phylip-3.697 in order to elucidate the phylogenetic relationship from a genome-wide perspective. The population genetic structure was investigated using the program ADMIXTURE v1.3.0. Assumed genetic clusters *K* was set 2 to 7, with 10,000 iterations per run. GCTA v1.91.5 (J. Yang, Lee, Goddard, & Visscher, 2013) was used to perform PCA to evaluate the population genetic structure.

### Population genetics analysis

To identify candidate regions potentially affected by selection, the fixation statistic (F_ST_) and nucleotide diversity (π) were calculated by VCFtools v0.1.13 (Danecek et al., 2011), with sliding windows of 50 kb and a 5-kb overlap between adjacent windows.

### RNA library construction and sequencing

Total RNA was extracted from the leaves of 12 samples using the RNAprep Pure Plant Kit (Tiangen DP441), and genomic DNA contamination was removed using the DNase I (Tiangen). Then, the isolated RNA was used for cDNA library construction using the NEBNext Ultra RNA Library Preparation Kit for Illumina (New England Biolabs, Ipswich, MA, USA), with a fragment length of approximately 150 bp. The cDNA library was paired-end sequenced on the Illumina NovaSeq 6000 platform.

### mRNA analysis

Mapping filtered reads to the reference genome of *S. pohuashanensis* using Hisat2 software.. Read count and the level of gene expression were quantified by the featureCounts program(Liao, Smyth, & Shi, 2014). The differential expression genes were then measured using the DESeq2 program(Love, Huber, & Anders, 2014), with the following criteria: FDR < 0.01 and absolute foldchange >1.

## Results

### Genome assembly and annotation

We generated PacBio HiFi sequences (40× coverage), Illumina short reads (90× coverage), and Hi-C sequences (100× coverage) for *S. pohuashanensis* (Hance) Hedl. We used the K-mer method to evaluate the *S. pohuashanensis* genome size based on Illumina short reads, and the estimated genome size was ∼668 Mb (Supplementary Fig. 1). Using the long reads, we assembled 426 contigs and generated a *S. pohuashanensis* genome of 660 Mb with a contig N50 of 28.2 Mb (Table 1); the corresponding values for the related species *M. domestica* cv. Gala are 665 Mb and 2.3 Mb, respectively, indicating that *S. pohuashanensis* and the domesticated apple have similar genome sizes.

**Table 1.** Assembly statistics of *S. pohuashanensis* genome.

| Category | *S. pohuashanensis* |  |  |  |  |
| --- | --- | --- | --- | --- | --- |
|  | Numbers | N50<br>(Mb) | Longest<br>(Mb) | Size<br>(Mb) | Percentage of assembly |
| Contigs | 426 | 28.2 | 43.7 | 660.1 | 100 |
| Anchored | 123 | 36.7 | 49.4 | 641.0 | 97.1 |
| Anchored and oriented | 39 | 30.6 | 43.7 | 623.3 | 97.2 |
| Gene annotated | 44,741 | NA | NA | 171.3 | 25.9 |
| Repeat sequence | NA | NA | NA | 451.8 | 68.4 |

With the aid of Hi-C data, the 426 contigs were clustered into 17 genetic groups, indicating that *S. pohuashanensis* has 17 chromosomes. We anchored ∼641 Mb of sequence onto 17 pseudochromosomes, of which 97.2% could be oriented. As expected, we found that the interaction intensities of the diagonal regions were stronger than those of the off-diagonal regions, indicating that the contigs were well positioned on the pseudochromosomes (Supplementary Fig. 2).

We next evaluated the assembly completeness using different methods. First, we mapped the Illumina reads and found that more than 98.60% were properly mapped to the new assembly (Supplementary Table 1). More than 97% of 1440 highly conserved embryophyte genes were identified as complete BUSCOs (Supplementary Table 2). The whole-genome high long terminal repeat (LTR) Assembly Index (LAI) (Ou, Chen, & Jiang, 2018) score is also an important indicator for the genome assembly quality. The LAI score for our assembly was 17.81, indicated that the assembly quality of *S. pohuashanensis* has reached the level of reference genome (Fig. 2D). As the most genomes, the *S. pohuashanensis* sequences positioned near the telomere were enriched of coding genes while having a lower-than-average level of repeat sequences. And the pericentromeric regions were enriched for repeat sequences but showed a deficit for coding genes compared to the genome-wide average (Fig. 2A, B).

**Fig. 2.**
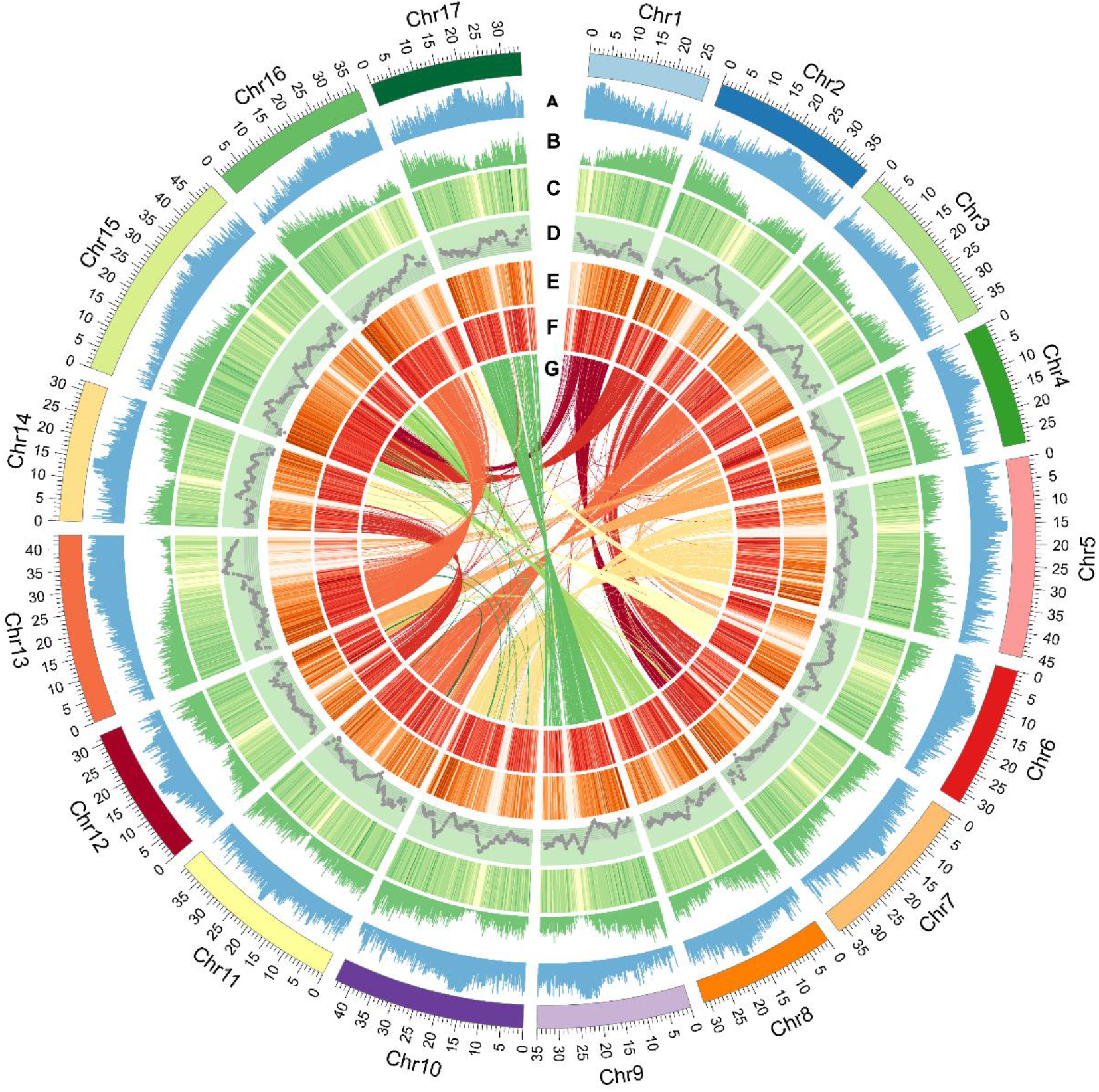
High-quality assembly of 17 chromosomes. **A**. Repeat sequence density (window size 200 kb). **B**. Gene density (window size of 200 kb). **C**. Gene expression level (FPKM value) in the control group (window size of 200 kb). **D**. LAI score (window size of 3 Mb, slide of 300 kb). Score: 0<LAI<10, Draft: 10≤LAI<20, Reference: LAI≥20, Gold. The boundary value of the dark green block of the LAI color block from the inside to the outside of the circle center is 10 and 20 respectively. Most LAI scores are >10, which indicates that the quality of the *Sorbus pohuashanensi* genome assembly is relatively high. **E**. SNP density (window size of 300 kb). **F**. Indel density (window size of 300 kb). **G**. Relationship between syntenic blocks.

A total of 46,870 protein-coding genes corresponding to 113,718 transcripts were identified, among which more than 95% could be annotated using at least one of the following protein-related databases: the GO database (45.56%), the KEGG database (32.77%), the KOG database (51.67%), the TrEMBL database (92.53%), and the NR database (95.36%), indicating the gene predictions were of high quality. In addition, we predicted 2822 tRNAs, 4863 rRNAs, and 111 miRNAs (Supplementary Table 1).

Repetitive elements are major components of genomes and one of the major causes for genome expansion and species divergence (W. Zhou, Liang, Molloy, & Jones, 2020). Approximately 68.44% of the assembly sequences were annotated as repetitive elements (Supplementary Table 3), which is higher than the percentages in domesticated apple (58.72%) and wild apples (56.93%–59.40%) (Sun et al., 2020).

### Gene family expansion and contraction

We selected nine sequenced species, namely *Amborella trichopoda, V. vinifera, Arabidopsis thaliana, Gossypium hirsutum, Populus trichocarpa, Fragaria vesca, Prunus persica, P. communis*, and *M. domestica*, together with *S. pohuashanensis* to analyze gene expansion and contraction. We used Orthofinder to cluster the protein sequences into families and found 4520 families that were shared by all ten species (Fig. 3A). We found 919 *S. pohuashanensis*-specific families that were composed of 3564 genes; these families were enriched in several pathways involved in the LTRs expansion such as DNA integration, RNA-dependent DNA biosynthetic process, and DNA recombination, indicating that LTRs play important roles in species-specific gene formation. Moreover, we also found that species-specific genes were enriched in photosynthesis-related pathways in both GO and KEGG analysis. Given that *S. pohuashanensis* plants grow on half sunny slopes, shady slopes, and valley environments, which are not conducive to photosynthesis, photosynthesis in *S. pohuashanensis* may have been enhanced by the retention of photosynthesis-related genes.

**Fig. 3.**
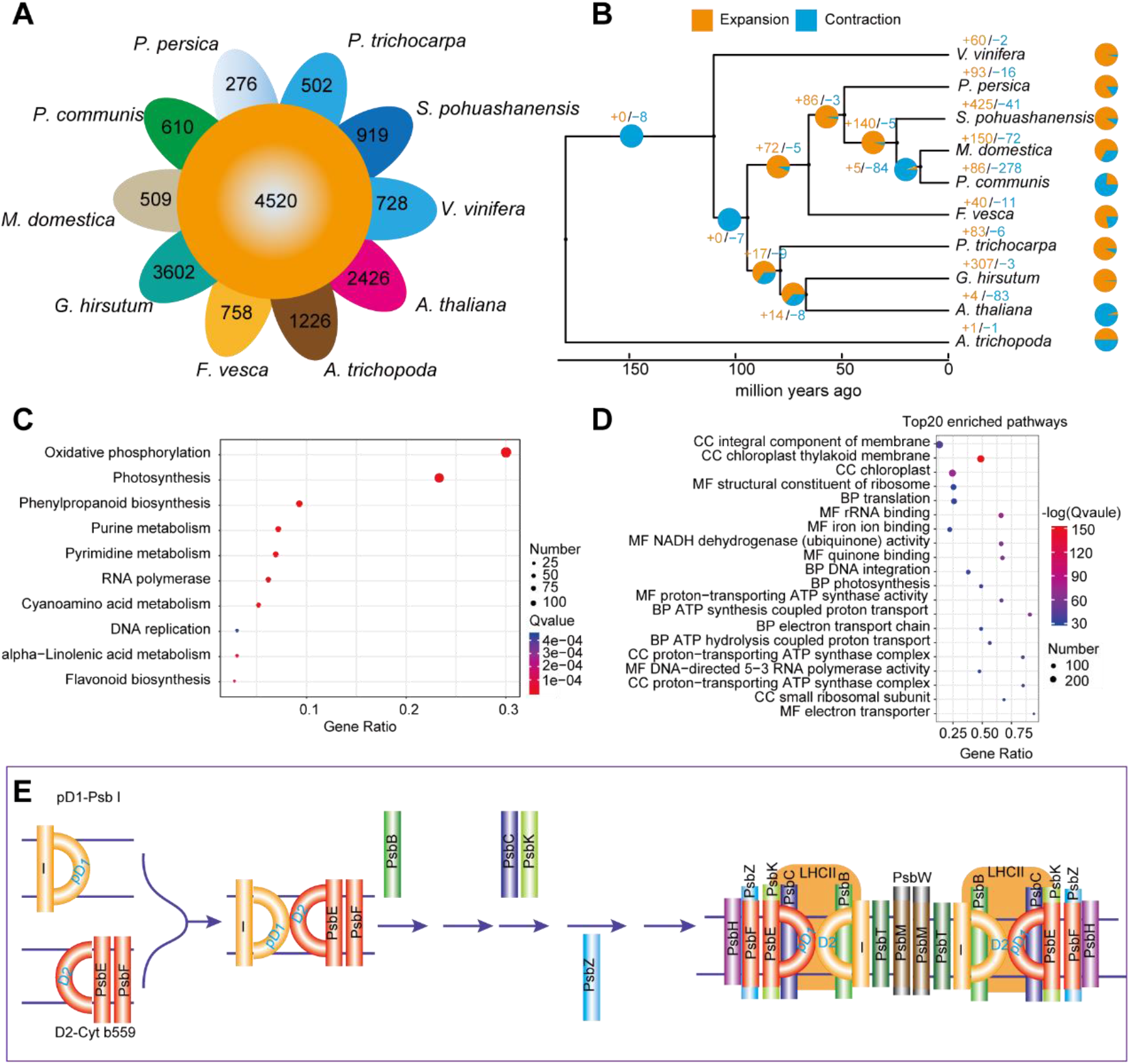
Evolution and low-temperature adaptation of the *Sorbus pohuashanensi* genome. **A**. A Venn diagram of specific and shared orthologs among ten species (*Vitis vinifera, Prunus persica, S. pohuashanensi, Malus domestica, Pyrus communis, Fragaria vesca, Populus trichocarpa, Gossypium hirsutum, Arabidopsis thaliana*, and *Amborella trichopoda*), identified based on gene family cluster analysis. Each number in the diagram represents the number of gene families within a group. **B**. Expansion and contraction of gene families. **C**. KEGG pathway enrichment analysis of expanded gene families. **D**. Top 20 enriched GO pathways. **E**. Examples of genes from expanded gene families involved in photosystem II (PSII) assembly. The genes shown are involved in the following steps: assembly of the precursor D1-PsbI (pD1-PsbI) and D2-cytochrome b559 (D2-Cyt b559) precomplexes, assembly of the reaction-center complex (PsbB/CP47), incorporation of PsbC (CP43) and PsbK to form the OEC-less PSII core monomer, and assembly of PsbZ to form the PSII core monomer.

Using 1241 single-copy genes, we constructed a phylogenetic tree and used the MCMCTREE program in PAML to estimate the divergence time. Based on the time tree, we used Computational Analysis of gene Family Evolution (CAFE) to estimate the number of gene families that experienced expansion or contraction. For all species except for *A. trichopoda, A. thaliana*, and *P. communis*, the number of gene families that experienced expansion was larger than the number that experienced contraction. We found that 425 gene families (4180 genes) and 41 gene families (82 genes) have expanded and contracted in *S. pohuashanensis*, respectively (Fig. 3B).

KEGG enrichment analysis showed that the expanded families were significantly enriched in photosynthesis pathways, including oxidative phosphorylation, photosynthesis, phenylpropanoid biosynthesis, and flavonoid biosynthesis (Fig. 3C). GO enrichment analysis also showed that the photosynthesis-related pathways chloroplast thylakoid membrane and photosynthesis were among the top 20 enriched pathways (Fig. 3D). The gene families encoding core components of photosystem II (PSII), including PSII protein D1, PSII protein D2, PsbB, PsbC, PsbK, and PsbZ, and ATP synthase were expanded in *S. pohuashanensis*, indicating that expansion of the photosynthesis pathway may have helped *S. pohuashanensis* to generate more energy to adapt to high-altitude and low-temperature environments (Fig. 3E).

### Evolution of the *S. pohuashanensis* genome

Whole-genome duplications (WGDs) have played important roles in the evolutionary history of angiosperms. Previous studies suggested that one to two genome duplications preceded angiosperm diversification, and only one known angiosperm, *A. trichopoda*, did not experience additional WGDs (Defoort, Van de Peer, & Carretero-Paulet, 2019). We first analyzed the syntenic blocks between *S. pohuashanensis* and *A. trichopoda* and found that one gene in *A. trichopoda* corresponded to an average of two orthologs in *S. pohuashanensis*, indicating that a WGD occurred in the *S. pohuashanensis* lineage. For example, we found that a CBL-interacting protein kinase 12 (evm_27.model.AmTr_v1.0_scaffold00007.215) from the *A. trichopoda* has four orthologs in both *S. pohuashanensis* and *M. domestica* (Supplementary Fig. 6B). Next, we used MCScanX to identify the syntenic blocks within *S. pohuashanensis*, and found a large number of syntenic blocks within this species (Supplementary Fig. 6A, Fig. 2G). The chromosome number in Rosaceae is 7, 8, or 9; for example, *F. vesca* has 7 chromosomes and *P. persica* has 8 chromosomes (Shulaev et al., 2008). Both of these species only experienced the core-eudicot γ whole-genome triplication (WGT) (Qiao et al., 2019). Both the synonymous substitution rate (Ks) and 4DTv value distributions showed that *S. pohuashanensis* experienced two rounds of WGD: one is the core eudicot γ WGT event (the Ks peak is 2.59 and the 4DTv peak is 0.0575) and the other is a recent WGD (the Ks peak is 0.18 and 4DTv peak is 0.36) (Supplementary Fig. 3). This finding is further supported by the clear-cut chromosomal DNA fragmentation patterns of *S. pohuashanensis* when compared with those of *P. persica and F. vesca* (Supplementary Fig. 6D).

*P. communis, M. domestica*, and *S pohuashanensis* all have 17 chromosomes, and there are conserved syntenic relationships among all orthologous chromosomes (Supplementary Fig. 6). We next evaluated the species divergence times (Supplementary Fig. 3) and found that *S. pohuashanensis* and *M. domestica* separated from the common ancestor 5.18 MYA, and the corresponding values for *P. communis, P. persica*, and *F. vesca* were 6.52 MYA, 46.4 MYA, and 83.1 MYA, respectively. We also calculated the time of the recent WGD for *S. pohuashanensis* to be 26.0 MYA, which is earlier than the time of *S. pohuashanensis* and *M. domestica* divergence but later than the time of *S. pohuashanensis* and *P. persica* divergence, supporting the notion that the recent WGD occurred in the common ancestor of *P. communis, M. domestica*, and *S. pohuashanensis* when diverged from *P. persica*.

Through whole chromosome alignments, we found that a large number of syntenic blocks among the non-homologous chromosomes, e.g. Chr01 and Chr07, Chr03 and Ch11, Chr04 and Chr12, Chr05 and Chr10, Chr08 and Chr17, and Chr13 and Chr16 were collinear, further supporting the existence of a recent WGD. We also compared the LTR insertion time among five species from Rosaceae (*F. vesca, P. persica, P. communis, M. domestica* and *S. pohuashanensis*) and found that LTR bursts mainly occurred between 0–1.25 MYA in *P. communis* (peak at 0.20 MYA) and *M. domestica* (peak at 0.25 MYA), and between 0–2 MYA in *P. persica* (peak at 0.39 MYA). There was a longer period of LTR insertion (between 0–4 MYA) in *F. vesca* (peak at 0.32 MYA). In contrast, an LTR burst in *Sorbus pohuashanensis* mainly occurred between 0–0.75 MYA with a peak at 0.11 MYA, indicating that the LTRs in *S. pohuashanensis* were more active than those in the other four species (Supplementary Fig. 3C).

### Dynamic changes of duplicated genes

Gene duplication play an important role in genome evolution. A total of 43,633 out of 46,870 genes were identified as duplicate genes, including 25,604 derived from WGD (58.7%), 2903 from tandem duplication (TD, 6.7%), 3462 from proximal duplication (PD, 7.9%), 5778 from transposed duplication (TRD, 13.2%), and 5886 from dispersed duplication (DSD, 13.5%) (Supplementary Table 4). The *Ks* (number of substitutions per synonymous site), *Ka* (number of substitutions per nonsynonymous site), and *Ka*/*Ks* ratio were calculated for different modes of duplication. We found that TD and PD genes had qualitatively higher *Ka*/*Ks* ratios than genes derived from other modes of duplication (Supplementary Fig. 4). The TD and PD gene pairs had relatively smaller *Ks* values (Supplementary Fig. 4). This finding supported a previous finding that younger TD and PD genes that have been preserved have experienced more rapid sequence divergence than other gene classes (Supplementary Fig. 4) (Qiao et al., 2019). Compared with all genes, the TD (439 out of 4,212) and PD (670 out of 4212) genes were enriched in genes from expanded gene families (Fisher test, P < 2.16 × 10^−16^), indicating that TD and PD were two of the major mechanisms for gene family expansion. KEGG analysis showed that the TD and PD genes from expanded gene families were enriched in phenylpropanoid biosynthesis, cyanoamino acid metabolism, plant-pathogen interaction, and flavonoid biosynthesis, suggesting that the TD and PD genes in *S. pohuashanensis* play important roles in environmental stress tolerance (Supplementary Fig. 5).

### Genomic divergence between simple-leaf and compound-leaf species

We collected 22 *Sorbus* L. samples from 11 species that are mainly from China: seven species with compound leaves, and four with simple leaves (Supplementary Table 5). Approximately 180 Gb of Illumina short reads were generated using the Illumina NovaSeq 6000 platform, and the average coverage depth for each sample was ∼10×. The BWA and GATK pipeline was used to map short reads to the new *S. pohuashanensis* reference genome and identify SNPs. On average, 95.46% of the reads for each accession were successfully aligned (Supplementary Table 5). We identified 8,770,043 high-quality SNPs and 912,343 Indels, with an average of 13.2 SNPs and 1.46 Indels per kb; these numbers were higher than those previously reported for diploid cotton (Ma et al., 2018), indicating that *Sorbus* L has high genomic diversity. Among the SNPs, 2,417,482 were located within protein-coding genes, 1,497,959 were located in upstream or downstream regions, and the remaining 4,854,602 SNPs were located in intergenic regions (Fig. 2E, F). In the coding regions, we annotated 378,589 (4.3% of the total) nonsynonymous SNPs and 303,424 synonymous, 926 stop-loss, and 6,907 stop-gain SNPs that caused amino acid changes, elongated transcripts, or premature stops (Supplementary Table 7).

We next analyzed the phylogenetic relationships among the 22 samples. The phylogenetic tree showed that the four simple-leaf species (a group named G1) clustered together, and the compound-leaf species (a group named G2) clustered together (Fig. 4A). Principal component analysis (PCA) (Fig. 4B) further supported the phylogenic results and showed that the G2 members have a closer phylogenetic relationship than those in G1. The nucleotide diversity (π) values for G1 and G2 were 3.2×10^−3^ and 2.5 ×10^−3^, respectively (Fig. 4A). We next used ADMIXTURE to analyze the population structure with the hypothesized number of populations ranging from *K* = 2 to *K* = 7 (Fig. 4C). We found that when *K* = 2, the compound-leaf species (G2) and simple-leaf species (G1) could be well separated from each other. As K increased from 3 to 5, the G2 group was divided into subgroups that perfectly matched the phylogenetic relationship (Fig. 4A).

**Fig. 4.**
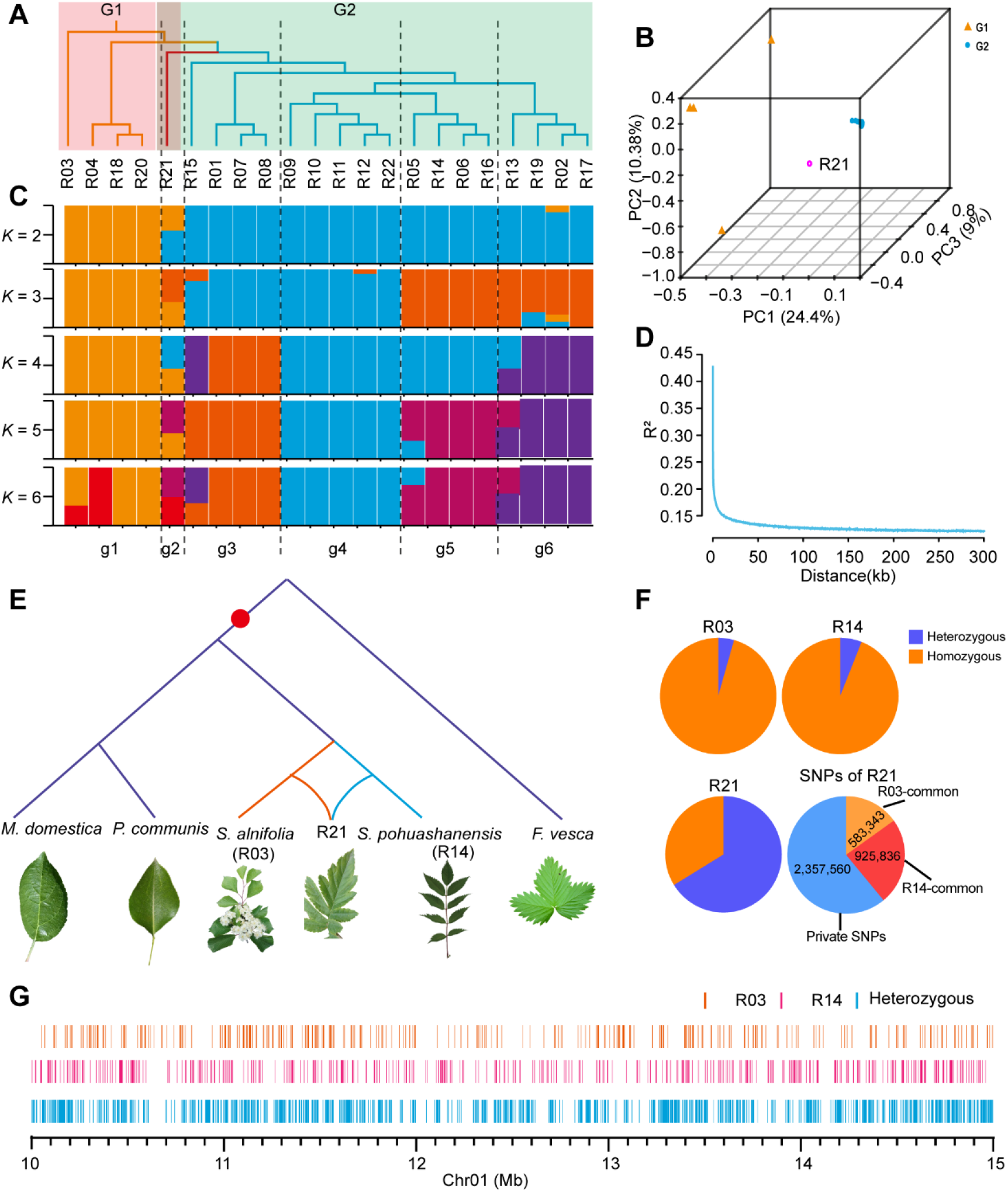
Resequencing and analysis of population structure and evolutionary relationships. **A**. Phylogenetic tree of 22 resequencing samples. **B**. Principal component analysis (PCA) of G1 (group1) and G2 (group2). **C**. Population structure (*K* = 2 to 6). **D**. Rate of linkage disequilibrium (LD) decay (R^2^). **E**. The suggested evolutionary relationship among R21, R03 and R14. **F**. Proportion of heterozygous SNPs in R03, R14, and R21. **G**. Genotype composition of R21 on Chr01from 10-15Mb. Orange lines represent the genotype of R21 is same with the R03, pink lines represent the genotype of R21 is same with the R14, and blue lines represent the heterozygous genotypes.

We next checked the geographical origin of the tested samples. The three species introduced from Europe (R19, R02, and R17) may have originated from *S. pohuashanensis* R13 (Haerbin, Heilongjiang, China, 126°34’E, 45°44′N). *S. pohuashanensis* R07 (Pangquan Ditch, 111°29’E, 37°50’N) and R08 (Tuoliang Mountain, 113°49’E, 38°43’N) clustered together with *S. discolor* R01 (Baihua Mountain, 117°57’E, 35°34’N), and they may have originated from *S. hupehensis* Schneid R15 (from Wufeng County, Hubei, China, 111°2’E, 30°11’N). The four compound from Shandong Province, *S. pohuashanensis* varieties R05, R06 and R14 and *S. discolor* var. *paucijuga* D.K.Zang et P.C.Huang R16, clustered together; *S. discolor* var. paucijuga possibly originated from *S. pohuashanensis*. The five *S. pohuashanensis* sample plants (R09, R10, R11, and R12) from North China have a very close relationship. These results suggested that the phylogenetic relationships among the compound-leaf samples were associated with their geographic locations.

We next calculated the fixation index (F_*ST*_) value between G1 and G2 to analyze the population differentiation. The results showed that G1 and G2 were significantly divergent with an average F_*ST*_ value equal to 0.5216. The highest 5% of the F_*ST*_ value was used as the threshold to select the gene sweep region. The region was 86.64 Mb containing 5484 genes. These genes were enriched in several pathways including catalytic activity, binding, transporter activity, and nucleic acid binding transcription factor activity. As there were only four simple-leaf samples in G1, we only calculated the linkage disequilibrium (LD) decay value for G2 and found that the pairwise correlation coefficient (R^2^) decreased from the maximum value (0.4714) to the half at 0.22 kb, the distance was much smaller than that of domesticated crops such as rice, soybean, and cotton (Fig. 4D) (Huang et al., 2010; Z. Zhou et al., 2015).

### Natural hybridization between simple-leaf species and compound-leaf species

It is very interesting that the R21 has a variant leaf shape (Fig. 4E) that is a cross between the simple-leaf and compound-leaf type. Although it is very difficult to create hybrids between simple-leaf species and compound-leaf species through artificial means, we predicted that R21 may have been generated by natural hybridization between a simple-leaf species and compound-leaf species because it coexists with the compound-leaf species *S. pohuashanensis* R14 and the simple-leaf species *S. alnifolia* (Sieb. et Zucc.) K. Koch R03 on Kunyun Mountain (121°44′E, 37°17′E,) (Fig. 4E). To test this hypothesis, we used R03 and R14 as the possible parents of R21 to analyze the SNPs. A total of 3,866,739 SNPs were identified among R03, R14, and R21. Then we analyzed the SNP heterozygosity and found that the SNPs heterozygosities of R03, R14, and R21 were 4.4%, 6.0%, and 66.2%, respectively. Thus, it is clear that the genome of R21 is highly heterozygous. Then we analyzed the common and private SNPs in R21 and found that R21 has 583,343 SNPs in common with R03 and 925,836 in common with R14 (Fig. 4F, G). Compared with common SNPs, the number of private SNPs in R21 was much higher (2,357,560) and almost all the private SNPs were heterozygous. Based on the above results, we conclude that there are natural hybridization occurs between the simple-leaf and compound-leaf species.

### Pathways and genes involved in sunburn response

Sunburn is one of the most serious types of damage faced by *S. pohuashanensis*. Here we used mRNA-seq to reveal the possible mechanism causing sunburn. We found that when *S. pohuashanensis* was grown under sheltering conditions, leaf color was uniform with a dark green color. Then we exposed *S. pohuashanensis* plants to sunlight. After 2.5 hours of sunlight treatment (hst), no visible change in leaf color was observed. However, at 26.5 hst, the leaves were brown, and at 146.5 hst necrotic spots were observed on the leaves (Fig. 5A, Supplementary Table 6). We took samples at 0, 2.5, 26.5, and 146.5 hst for mRNA-seq; these samples were designated CK, L1, L2 and L3, respectively. A total of 934, 1170, and 1022 up-regulated genes were identified from comparisons between L1 and CK, L2 and CK, and L3 and CK, respectively. The corresponding values for down-regulated genes were 1775, 1764, and 2011, respectively. Thus, it is clear that the number of down-regulated genes is much higher than that of up-regulated genes, suggesting that gene transcription was restricted under sunlight.

**Fig. 5.**
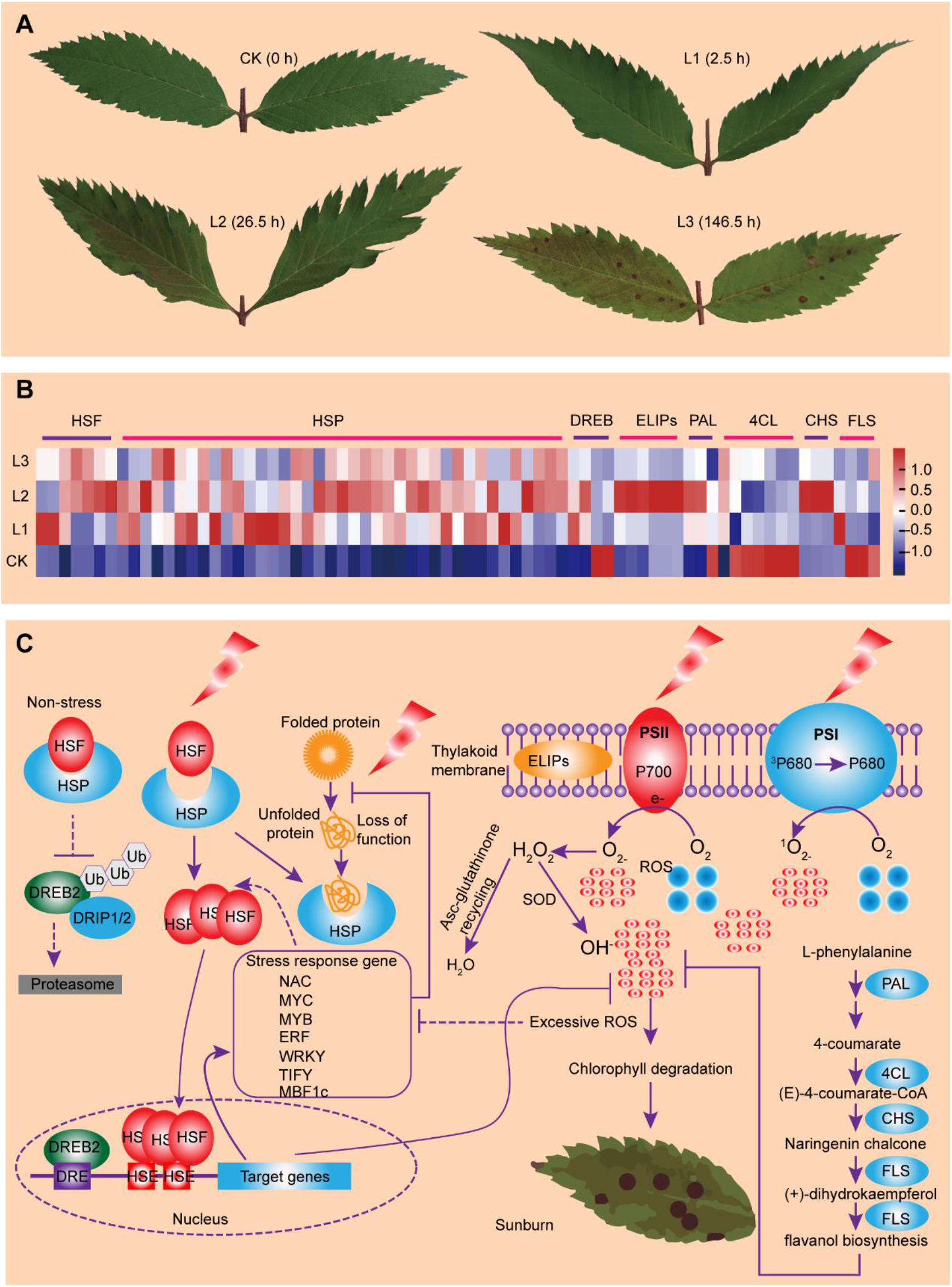
Sunburn phenotype and response pathway. **A**. Leaf burn phenotypes after different treatment periods. **B**. Heatmap of the expression levels of the core sunburn response genes after different periods of sunlight. **C**. Sunburn response pathway. Upon sunlight stress, the HSP and HSF pathways are activated. The high number of misfolded proteins triggers the recruitment of HSPs to their clients. The free HSFs can bind to the HSE elements of target genes and activate signaling pathways to remove reactive oxygen species (ROS). Genes of the flavanol biosynthesis pathway showed differential responses to sunlight treatments, with PAL, 4CL, CHS, and FLS being up-regulated. Flavanol together with enzymatic antioxidants help to eliminate the ROS. MYC, MYB, ERF, WRKY and TIFY TFs were down-regulated, indicating that their activities may be restricted by the excessive ROS levels induced by high temperature and solar radiation. The rate of ROS generation exceeds the rate of its elimination, which leads to the failure to remove the ROS, resulting in chlorophyll degradation and eventually leading to mesophyll tissue necrosis.

We identified 182 genes including early-light induced proteins (ELIPs), heat shock proteins (HSPs), heat shock factors (HSFs), and downstream transcription factors (TFs) (e.g. DREBs, ERFs, and WRKYs) potentially involved in high-temperature response (Fig. 5B, Supplementary Table 8). HSPs and HSFs were up-regulated at L1, L2, and L3, indicating that the HSP and HSF pathways were activated. Upon sunlight stress, a high number of misfolded proteins accumulate, which triggers the recruitment of HSPs to their client HSFs. After being freed by HSPs, HSFs can bind to the HSE elements of target genes and activate signaling pathways to remove various reactive oxygen species (ROS). For example, flavanol together with enzymatic antioxidants help to eliminate ROS. Consistent with this, flavanol biosynthesis pathway genes showed differential expression in response to sunlight treatments, with PAL, 4CL, CHS, and FLS being up-regulated (Fig. 5C).

Interestingly, most of the differentially expressed TFs including MYCs, MYBs, ERFs, WRKYs and TIFYs were down-regulated, indicating that their activities may be restricted by the excessive production of ROS induced by high temperature and solar radiation. As the rate of ROS generation exceeds the rate of degradation, there is a failure to remove the ROS, resulting in chlorophyll degradation and eventually leading to mesophyll tissue necrosis (Fig. 5C).

## Discussion

Here we performed de novo assembly of the genome of *S. pohuashanensis* by integrating data from Pacbio HiFi sequencing and Hi-C, providing the first reference genome for the *Sorbus* genus. The new reference genome will both contribute to *Sorbus* species classification and basic research. We also sequenced 11 species of *Sorbus* and found that compound-leaf species have a distant phylogenetic relationship with the simple-leaf species. Although only four simple-leaf species were used in this study, we found that they could be separated from each other in PCA analysis. In contrast, we found that the samples with compound leaves clustered together in PCA analysis and had a lower nucleotide diversity. *M. domestica*, and *P. communis* have simple leaves; thus, it is clear that simple-leaf *Sorbus* species were formed earlier than the compound-leaf *Sorbus* species, and that compound-leaf species may originate from a simple-leaf species. *S. pohuashanensis* and six other compound-leaf species were used in the phylogenetic analysis, but we found *S. pohuashanensis* did not cluster together in the phylogenetic tree. One possible reason for this is that it is very easy to make mistakes in species classification using only shape and geographic origin. For example, R16 (*S. discolor* var. paucijuga D. K. Zang et P. C. Huang) is the sister species of R06 and R14 (*Sorbus pohuashanensis*) in the phylogenetic tree, and it clustered together with R06 and R14 in PCA, indicating that they were from the same species. Similarly, the R01 (*S. discolor* (Maxim.) Maxim) has a very close relationship to R06 and R14 (*S. pohuashanensis*), and they should be considered as the same species.

Natural hybridization between closely related species is common in plants, such as cotton and wheat, and is an important source of new species. Both the simple-leaf species *S. alnifolia* and the compound-leaf species *S. pohuashanensis* are located on Kunyu Mountain (Shandong Province, China), which raises the possibility for hybridization between these species. The existence of R21 provides strong evidence for the natural hybridization between simple-leaf and compound-leaf species. However, we found that only 66.2% of the SNPs in the genome were heterozygous, indicating that R21 is not from the F1 generation after hybridization, and is likely an offspring of an F1 hybrid.

Sunburn is a physiological disease in which damage occurs due to excessive heat and/or light radiation (visible light and ultraviolet light) (Munné-Bosch & Vincent, 2019). When the air temperature is maintained at a normal growth temperature, there may be a significant overlap between the excessive light response and the thermal stress response (Balfagón et al., 2019). During excessive light stress, the light reaction center becomes saturated, and the excess excitation energy may become harmful because it irreversibly destroys PSII (Murata, Takahashi, Nishiyama, & Allakhverdiev, 2007; Ruban, 2009, 2015). An imbalance between the rate of PSII destruction by light and the rate of PSII repair leads to photoinhibition, and the efficiency of photosynthesis continues to decline (Murata et al., 2007; Nishiyama, Allakhverdiev, & Murata, 2006). In addition to excessive light stress, heat stress also impairs photosynthesis by hindering the transport of PSII electrons due to an increase in thylakoid membrane fluidity, which causes the PSII light-harvesting complex (LHC) to fall off and reduces the integrity of PSII (Balfagón et al., 2019; Mathur, Agrawal, & Jajoo, 2014). The ELIPs, which accumulate in the thylakoid membrane, play important roles in photoprotection. When photoinhibition occurs, it has been described as an ELIP accumulation pattern, accompanied by a decrease in LHC levels due to D1 protein denaturation in PSII (Munné-Bosch & Vincent, 2019). We also found that six ELIPs were highly up-regulated and that the LHC type 1-like protein chlorophyll a-b binding protein was down-regulated after sunlight treatment, suggesting that ELIPs and the LHC play a role in photoprotection in *Sorbus pohuashanensis*.

Light intensities and heat stress induce the production of ROS via different mechanisms (Pospíšil, 2016). The production of ROS under stress conditions initially triggers a signal cascade, and if amounts sufficient to overwhelm the defense mechanism are produced, it will also cause cell damage. Singlet oxygen (^1^O_2_) is very harmful and may be the main ROS produced under excessive light (D’Alessandro & Havaux, 2019; Khorobrykh, Havurinne, Mattila, & Tyystjärvi, 2020). Short-term (24 h) high light treatment causes the accumulation of ^1^O_2(Shumbe et al., 2017)_; prolonged high light stress is therefore likely to cause the combined accumulation of ^1^O_2_ and H_2_O_2_ (Laloi & Havaux, 2015). After 24 hours of high heat and high light stress in *A. thaliana*, PSII function cannot be fully restored, and lipid peroxidation is usually the main event related to ROS production and oxidative stress in plants (Birtic et al., 2011). The reaction of ROS with lipids produces specific peroxides, especially lipid endoperoxides unique to ^1^O_2_ and lipid free radicals and lipid peroxyl free radicals, which are specific to the reduced oxygen form (Khorobrykh et al., 2020; Mano, Biswas, & Sugimoto, 2019). Some studies observed that the sunlit sides of plants contain greater amounts of flavonols than the shaded sides (Felicetti & Schrader, 2008). Flavonols are phenolic compounds that enhance light absorption from the UV and blue regions of the spectrum and act as scavengers of ROS molecules (Solovchenko & Merzlyak, 2008). We also found that key genes involved in flavonol biosynthesis pathways were up-regulated under both short-term (L1) and longer sunlight treatments (L2 and L3), suggesting that flavonols have conserved functions in ROS scavenging in *S. pohuashanensis*. HSP can bind to the unfolded proteins generated by sunlight stress, and HSF can bind to the HSE elements in the target heat-response genes. We found the that HSF-HSP pathway was activated at L1, L2 and L3, suggesting that HSF-HSP pathways play important roles in ROS scavenging.

In summary, we de novo assembled a very high-quality reference genome for the important *Sorbus* genus using technologically complementary sequencing technologies. Based on this reference-grade genome assembly, we comprehensively evaluated gene family expansion and contraction, genome evolution, population structure, and the mechanism underlying sunburn response to better understand *Sorbus* genetics and evolution. Of particular note, our resequencing data provided insights into natural hybridization between different *Sorbus* species. Our study also identified multiple networks underlying the sunburn response, which can promote efforts to apply *S. pohuashanensis* as a horticultural crop. Thus, our findings deepen the understanding of *Sorbus* population evolution, genome evolution, and sunburn response mechanisms, and provide an important resource for the introduction and breeding of *Sorbus* spp.

## Acknowledgments

We thank the National Forest Genetic Resources Platform (NFGRP) for providing the *S. pohuashanensis* plant resources. This work was supported by the Fund of the National Natural Science Foundation of China (grant number 31770369) to J. Z.

## Data availability

Raw data from this study were deposited in the NCBI SRA (Sequence Read Archive) database under the Bioproject ID: PRJNA716887. The assembled genome sequences and gene annotation have been deposited at FigShare, https:doi.org/10.6084/m9.figshare.14920695. The whole genome sequence data have been deposited in the Genome Warehouse in National Genomics Data Center, under accession number GWHBDNT00000000 that is publicly accessible at https://ngdc.cncb.ac.cn/gwh.

## Author contributions

D. Zhao, Y. Zhang and Y. Lu designed and performed the experiments; D. Zhao, Y. Zhang, Y. Lu, L. Fan and Z. Zhang analyzed the data; D. Zhao and J. Zheng wrote and revised the manuscript, respectively; J. Zheng and M. Chai supervised the research. All authors read and approved the final manuscript for publication.

## References

Altschul, S. F., Gish, W., Miller, W., Myers, E. W., & Lipman, D. J. (1990). Basic local alignment search tool. Journal of molecular biology, 215(3), 403–410.

Balfagón, D., Sengupta, S., Gómez-Cadenas, A., Fritschi, F. B., Azad, R. K., Mittler, R., & Zandalinas, S. I. (2019). Jasmonic Acid Is Required for Plant Acclimation to a Combination of High Light and Heat Stress. Plant Physiology, 181(4), 1668–1682. doi:10.1104/pp.19.00956

Birtic, S., Ksas, B., Genty, B., Mueller, M. J., Triantaphylidès, C., & Havaux, M. (2011). Using spontaneous photon emission to image lipid oxidation patterns in plant tissues. The Plant Journal, 67(6), 1103–1115.

Boeckmann, B., Bairoch, A., Apweiler, R., Blatter, M.-C., Estreicher, A., Gasteiger, E., … Phan, I. (2003). The SWISS-PROT protein knowledgebase and its supplement TrEMBL in 2003. Nucleic Acids Research, 31(1), 365–370.

Buchfink, B., Xie, C., & Huson, D. H. (2015). Fast and sensitive protein alignment using DIAMOND. Nature methods, 12(1), 59–60.

D’Alessandro, S., & Havaux, M. (2019). Sensing β-carotene oxidation in photosystem II to master plant stress tolerance. New Phytologist, 223(4), 1776–1783. doi:https://doi.org/10.1111/nph.15924

Danecek, P., Auton, A., Abecasis, G., Albers, C. A., Banks, E., DePristo, M. A., … Sherry, S. T. (2011). The variant call format and VCFtools. Bioinformatics, 27(15), 2156–2158.

Defoort, J., Van de Peer, Y., & Carretero-Paulet, L. (2019). The Evolution of Gene Duplicates in Angiosperms and the Impact of Protein–Protein Interactions and the Mechanism of Duplication. Genome Biology and Evolution, 11(8), 2292–2305. doi:10.1093/gbe/evz156

DePristo, M. A., Banks, E., Poplin, R., Garimella, K. V., Maguire, J. R., Hartl, C., … Daly, M. J. (2011). A framework for variation discovery and genotyping using next-generation DNA sequencing data. Nature genetics, 43(5), 491–498. doi:10.1038/ng.806

Dimmer, E. C., Huntley, R. P., Alam-Faruque, Y., Sawford, T., O’Donovan, C., Martin, M. J., … Eberhardt, R. (2012). The UniProt-GO annotation database in 2011. Nucleic Acids Research, 40(D1), D565–D570.

Du, H., Yu, Y., Ma, Y., Gao, Q., Cao, Y., Chen, Z., … Liang, C. (2017). Sequencing and de novo assembly of a near complete indica rice genome. Nature Communications, 8(1), 1–12. doi:10.1038/ncomms15324

El-Tanany, M., Kheder, A., & Abdallah, H. R. (2019). Effect of some treatments on reducing sunburn in Balady Mandarin fruit trees (Citrus reticulata, Blanco). Middle East J. Agric. Res, 8(3), 889–897.

Ellinghaus, D., Kurtz, S., & Willhoeft, U. (2008). LTRharvest, an efficient and flexible software for de novo detection of LTR retrotransposons. BMC bioinformatics, 9(1), 18. doi:10.1186/1471-2105-9-18

Emms, D. M., & Kelly, S. (2019). OrthoFinder: phylogenetic orthology inference for comparative genomics. Genome Biology, 20(1), 238. doi:10.1186/s13059-019-1832-y

Felicetti, D. A., & Schrader, L. E. (2008). Photooxidative sunburn of apples: Characterization of a third type of apple sunburn. International journal of fruit science, 8(3), 160–172.

Fu, A., Wang, Q., Mu, J., Ma, L., Wen, C., Zhao, X., … Zuo, J. (2021). Combined genomic, transcriptomic, and metabolomic analyses provide insights into chayote (Sechium edule) evolution and fruit development. Horticulture Research, 8(1), 35. doi:10.1038/s41438-021-00487-1

Goodwin, I., McClymont, L., Turpin, S., & Darbyshire, R. (2018). Effectiveness of netting in decreasing fruit surface temperature and sunburn damage of red-blushed pear. New Zealand Journal of Crop and Horticultural Science, 46(4), 334–345.

Han, M. V., Thomas, G. W. C., Lugo-Martinez, J., & Hahn, M. W. (2013). Estimating Gene Gain and Loss Rates in the Presence of Error in Genome Assembly and Annotation Using CAFE 3. Molecular Biology and Evolution, 30(8), 1987–1997. doi:10.1093/molbev/mst100

Hoede, C., Arnoux, S., Moisset, M., Chaumier, T., Inizan, O., Jamilloux, V., & Quesneville, H. (2014). PASTEC: an automatic transposable element classification tool. PloS one, 9(5), e91929.

Huang, X., Sang, T., Zhao, Q., Feng, Q., Zhao, Y., Li, C., … Li, M. (2010). Genome-wide association studies of 14 agronomic traits in rice landraces. Nature genetics, 42(11), 961–967.

Jiao, Y., Peluso, P., Shi, J., Liang, T., Stitzer, M. C., Wang, B., … Ware, D. (2017). Improved maize reference genome with single-molecule technologies. Nature, 546(7659), 524–527. doi:10.1038/nature22971

Jurka, J., Kapitonov, V. V., Pavlicek, A., Klonowski, P., Kohany, O., & Walichiewicz, J. (2005). Repbase Update, a database of eukaryotic repetitive elements. Cytogenetic and genome research, 110(1-4), 462–467.

Kalyaanamoorthy, S., Minh, B. Q., Wong, T. K., von Haeseler, A., & Jermiin, L. S. (2017). ModelFinder: fast model selection for accurate phylogenetic estimates. Nature methods, 14(6), 587–589.

Kanehisa, M., & Goto, S. (2000). KEGG: kyoto encyclopedia of genes and genomes. Nucleic Acids Research, 28(1), 27–30.

Khorobrykh, S., Havurinne, V., Mattila, H., & Tyystjärvi, E. (2020). Oxygen and ROS in Photosynthesis. Plants, 9(1), 91.

Koonin, E. V., Fedorova, N. D., Jackson, J. D., Jacobs, A. R., Krylov, D. M., Makarova, K. S., … Rao, B. S. (2004). A comprehensive evolutionary classification of proteins encoded in complete eukaryotic genomes. Genome Biology, 5(2), R7.

Laloi, C., & Havaux, M. (2015). Key players of singlet oxygen-induced cell death in plants. Frontiers in plant science, 6, 39.

Li, H., Handsaker, B., Wysoker, A., Fennell, T., Ruan, J., Homer, N., … Durbin, R. (2009). The sequence alignment/map format and SAMtools. Bioinformatics, 25(16), 2078–2079.

Liao, Y., Smyth, G. K., & Shi, W. (2014). featureCounts: an efficient general purpose program for assigning sequence reads to genomic features. Bioinformatics, 30(7), 923–930.

Love, M. I., Huber, W., & Anders, S. (2014). Moderated estimation of fold change and dispersion for RNA-seq data with DESeq2. Genome Biology, 15(12), 550. doi:10.1186/s13059-014-0550-8

Ma, Z., He, S., Wang, X., Sun, J., Zhang, Y., Zhang, G., … Du, X. (2018). Resequencing a core collection of upland cotton identifies genomic variation and loci influencing fiber quality and yield. Nature genetics, 50(6), 803–813. doi:10.1038/s41588-018-0119-7

Mano, J. i., Biswas, M., & Sugimoto, K. (2019). Reactive carbonyl species: a missing link in ROS signaling. Plants, 8(10), 391.

Marchler-Bauer, A., Lu, S., Anderson, J. B., Chitsaz, F., Derbyshire, M. K., DeWeese-Scott, C., … Gonzales, N. R. (2010). CDD: a Conserved Domain Database for the functional annotation of proteins. Nucleic Acids Research, 39(Suppl_1), D225–D229.

Mathur, S., Agrawal, D., & Jajoo, A. (2014). Photosynthesis: response to high temperature stress. Journal of Photochemistry and Photobiology B: Biology, 137, 116–126.

Mi, H., Muruganujan, A., Ebert, D., Huang, X., & Thomas, P. D. (2018). PANTHER version 14: more genomes, a new PANTHER GO-slim and improvements in enrichment analysis tools. Nucleic Acids Research, 47(D1), D419–D426. doi:10.1093/nar/gky1038

Munné-Bosch, S., & Vincent, C. (2019). Physiological Mechanisms Underlying Fruit Sunburn. Critical Reviews in Plant Sciences, 38(2), 140–157. doi:10.1080/07352689.2019.1613320

Murata, N., Takahashi, S., Nishiyama, Y., & Allakhverdiev, S. I. (2007). Photoinhibition of photosystem II under environmental stress. Biochimica et Biophysica Acta (BBA)-Bioenergetics, 1767(6), 414–421.

Nguyen, L.-T., Schmidt, H. A., von Haeseler, A., & Minh, B. Q. (2014). IQ-TREE: A Fast and Effective Stochastic Algorithm for Estimating Maximum-Likelihood Phylogenies. Molecular Biology and Evolution, 32(1), 268–274. doi:10.1093/molbev/msu300

Nishiyama, Y., Allakhverdiev, S. I., & Murata, N. (2006). A new paradigm for the action of reactive oxygen species in the photoinhibition of photosystem II. Biochimica et Biophysica Acta (BBA)-Bioenergetics, 1757(7), 742–749.

Ossowski, S., Schneeberger, K., Lucas-Lledó, J. I., Warthmann, N., Clark, R. M., Shaw, R. G., … Lynch, M. (2010). The rate and molecular spectrum of spontaneous mutations in Arabidopsis thaliana. science, 327(5961), 92–94.

Ou, S., Chen, J., & Jiang, N. (2018). Assessing genome assembly quality using the LTR Assembly Index (LAI). Nucleic Acids Research, 46(21), e126–e126. doi:10.1093/nar/gky730

Ou, S., & Jiang, N. (2018). LTR_retriever: A Highly Accurate and Sensitive Program for Identification of Long Terminal Repeat Retrotransposons. Plant Physiology, 176(2), 1410–1422. doi:10.1104/pp.17.01310

Pospíšil, P. (2016). Production of reactive oxygen species by photosystem II as a response to light and temperature stress. Frontiers in plant science, 7, 1950.

Price, A. L., Jones, N. C., & Pevzner, P. A. (2005). De novo identification of repeat families in large genomes. Bioinformatics, 21(suppl_1), i351–i358.

Puttick, M. N. (2019). MCMCtreeR: functions to prepare MCMCtree analyses and visualize posterior ages on trees. Bioinformatics, 35(24), 5321–5322.

Qiao, X., Li, Q., Yin, H., Qi, K., Li, L., Wang, R., … Paterson, A. H. (2019). Gene duplication and evolution in recurring polyploidization–diploidization cycles in plants. Genome Biology, 20(1), 38. doi:10.1186/s13059-019-1650-2

Racsko, J., & Schrader, L. (2012). Sunburn of apple fruit: Historical background, recent advances and future perspectives. Critical Reviews in Plant Sciences, 31(6), 455–504.

Rice, P., Longden, I., & Bleasby, A. (2000). EMBOSS: the European Molecular Biology Open Software Suite. Trends in genetics : TIG, 16(6), 276–277. doi:10.1016/s0168-9525(00)02024-2

Ruban, A. V. (2009). Plants in light. Communicative & integrative biology, 2(1), 50–55.

Ruban, A. V. (2015). Evolution under the sun: optimizing light harvesting in photosynthesis. Journal of experimental botany, 66(1), 7–23.

Rustioni, L., Fracassetti, D., Prinsi, B., Geuna, F., Ancelotti, A., Fauda, V., … Failla, O. (2020). Oxidations in white grape (Vitis vinifera L.) skins: Comparison between ripening process and photooxidative sunburn symptoms. Plant Physiology and Biochemistry, 150, 270–278.

Shulaev, V., Korban, S. S., Sosinski, B., Abbott, A. G., Aldwinckle, H. S., Folta, K. M., … Veilleux, R. E. (2008). Multiple Models for Rosaceae Genomics. Plant Physiology, 147(3), 985–1003. doi:10.1104/pp.107.115618

Shumbe, L., d’Alessandro, S., Shao, N., Chevalier, A., Ksas, B., Bock, R., & Havaux, M. (2017). METHYLENE BLUE SENSITIVITY 1 (MBS1) is required for acclimation of Arabidopsis to singlet oxygen and acts downstream of β-cyclocitral. Plant, cell & environment, 40(2), 216–226.

Solovchenko, A., & Merzlyak, M. (2008). Screening of visible and UV radiation as a photoprotective mechanism in plants. Russian Journal of Plant Physiology, 55(6), 719–737.

Sun, X., Jiao, C., Schwaninger, H., Chao, C. T., Ma, Y., Duan, N., … Cheng, L. (2020). Phased diploid genome assemblies and pan-genomes provide insights into the genetic history of apple domestication. Nature genetics, 52(12), 1423–1432.

Tang, H., Krishnakumar, V., Li, J., & Zhang, X. (2015). jcvi: JCVI utility libraries. Zenodo.(doi: 10.5281/zenodo.31631).

Tarailo-Graovac, M., & Chen, N. (2009). Using RepeatMasker to identify repetitive elements in genomic sequences. Current protocols in bioinformatics, 25(1), 4.10.11-14.10.14.

Wang, D., Zhang, Y., Zhang, Z., Zhu, J., & Yu, J. (2010). KaKs_Calculator 2.0: a toolkit incorporating gamma-series methods and sliding window strategies. Genomics, proteomics & bioinformatics, 8(1), 77–80.

Wang, Y., Tang, H., DeBarry, J. D., Tan, X., Li, J., Wang, X., … Guo, H. (2012). MCScanX: a toolkit for detection and evolutionary analysis of gene synteny and collinearity. Nucleic Acids Research, 40(7), e49–e49.

Xu, Y., Bi, C., Wu, G., Wei, S., Dai, X., Yin, T., & Ye, N. (2016). VGSC: a web-based Vector Graph toolkit of genome Synteny and Collinearity. BioMed Research International, 2016.

Xu, Z., & Wang, H. (2007). LTR_FINDER: an efficient tool for the prediction of full-length LTR retrotransposons. Nucleic Acids Research, 35(Suppl_2), W265–W268. doi:10.1093/nar/gkm286

Yang, H., & Wang, K. (2015). Genomic variant annotation and prioritization with ANNOVAR and wANNOVAR. Nature protocols, 10(10), 1556–1566.

Yang, J., Lee, S. H., Goddard, M. E., & Visscher, P. M. (2013). Genome-wide complex trait analysis (GCTA): methods, data analyses, and interpretations. In Genome-wide association studies and genomic prediction (pp. 215–236): Springer.

Yang, Z. (1997). PAML: a program package for phylogenetic analysis by maximum likelihood. Computer applications in the biosciences, 13(5), 555–556.

Yang, Z., Ge, X., Yang, Z., Qin, W., Sun, G., Wang, Z., … Li, F. (2019). Extensive intraspecific gene order and gene structural variations in upland cotton cultivars. Nature Communications, 10(1), 1–13. doi:10.1038/s41467-019-10820-x

Yin, Y., Zhang, Y., Li, H., Zhao, Y., Cai, E., Zhu, H., … Liu, J. (2019). Triterpenoids from fruits of Sorbus pohuashanensis inhibit acetaminophen-induced acute liver injury in mice. Biomedicine & Pharmacotherapy, 109, 493–502.

Yu, G., Wang, L.-G., Han, Y., & He, Q.-Y. (2012). clusterProfiler: an R package for comparing biological themes among gene clusters. Omics: a journal of integrative biology, 16(5), 284–287. doi:doi.org/10.1089/omi.2011.0118

Yu, X., Wang, Z., Shu, Z., Li, Z., Ning, Y., Yun, K., … Liu, W. (2017). Effect and mechanism of Sorbus pohuashanensis (Hante) Hedl. flavonoids protect against arsenic trioxide-induced cardiotoxicity. Biomedicine & Pharmacotherapy, 88, 1–10.

Zhang, Z., Pei, X., Zhang, R., Lu, Y., Zheng, J., & Zheng, Y. (2020). Molecular characterization and expression analysis of small heat shock protein 17.3 gene from Sorbus pohuashanensis (Hance) Hedl. in response to abiotic stress. Molecular Biology Reports, 47(12), 9325–9335. doi:10.1007/s11033-020-06020-2

Zhang, Z., Xiao, J., Wu, J., Zhang, H., Liu, G., Wang, X., & Dai, L. (2012). ParaAT: a parallel tool for constructing multiple protein-coding DNA alignments. Biochemical and biophysical research communications, 419(4), 779–781.

Zhou, W., Liang, G., Molloy, P. L., & Jones, P. A. (2020). DNA methylation enables transposable element-driven genome expansion. Proc Natl Acad Sci U S A, 117(32), 19359–19366. doi:10.1073/pnas.1921719117

Zhou, Z., Jiang, Y., Wang, Z., Gou, Z., Lyu, J., Li, W., … Ma, Y. (2015). Resequencing 302 wild and cultivated accessions identifies genes related to domestication and improvement in soybean. Nature biotechnology, 33(4), 408–414.

Zwaenepoel, A., & Van de Peer, Y. (2018). wgd—simple command line tools for the analysis of ancient whole-genome duplications. Bioinformatics, 35(12), 2153–2155. doi:10.1093/bioinformatics/bty915

