## Supplementary material for "The genome sequence of *Sorbus pohuashanensis* provides insights into population evolution and leaf sunburn response": supply info

### Supplementary Information

#### Supplementary Figures

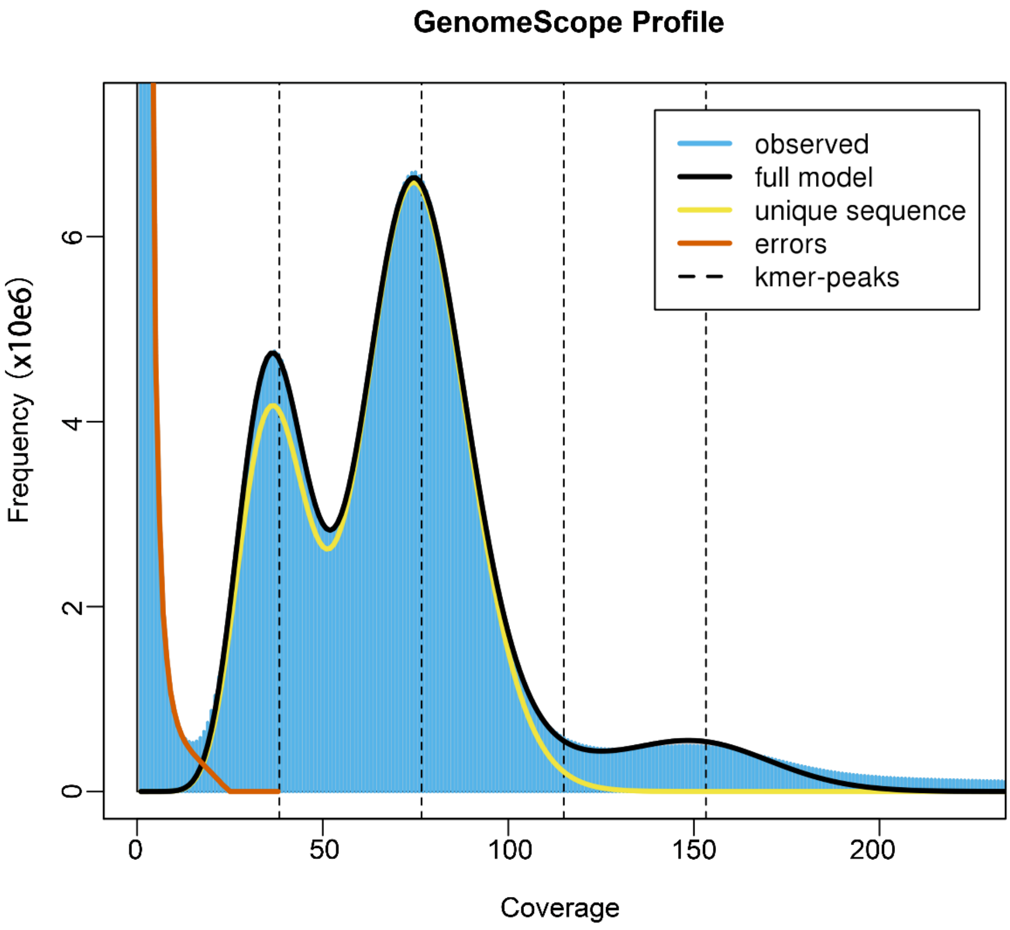


**Figure S1. The 19-mer distribution of Illumina short reads in S. pohuashanensis.** The x-axis shows the frequency or the number of times a given k-mer (k-mer depth). The y-axis shows the total number of k-mers with a given frequency (a given depth). Two peaks (blue line) were observed indicating heterozygosity in *S. pohuashanensis*.

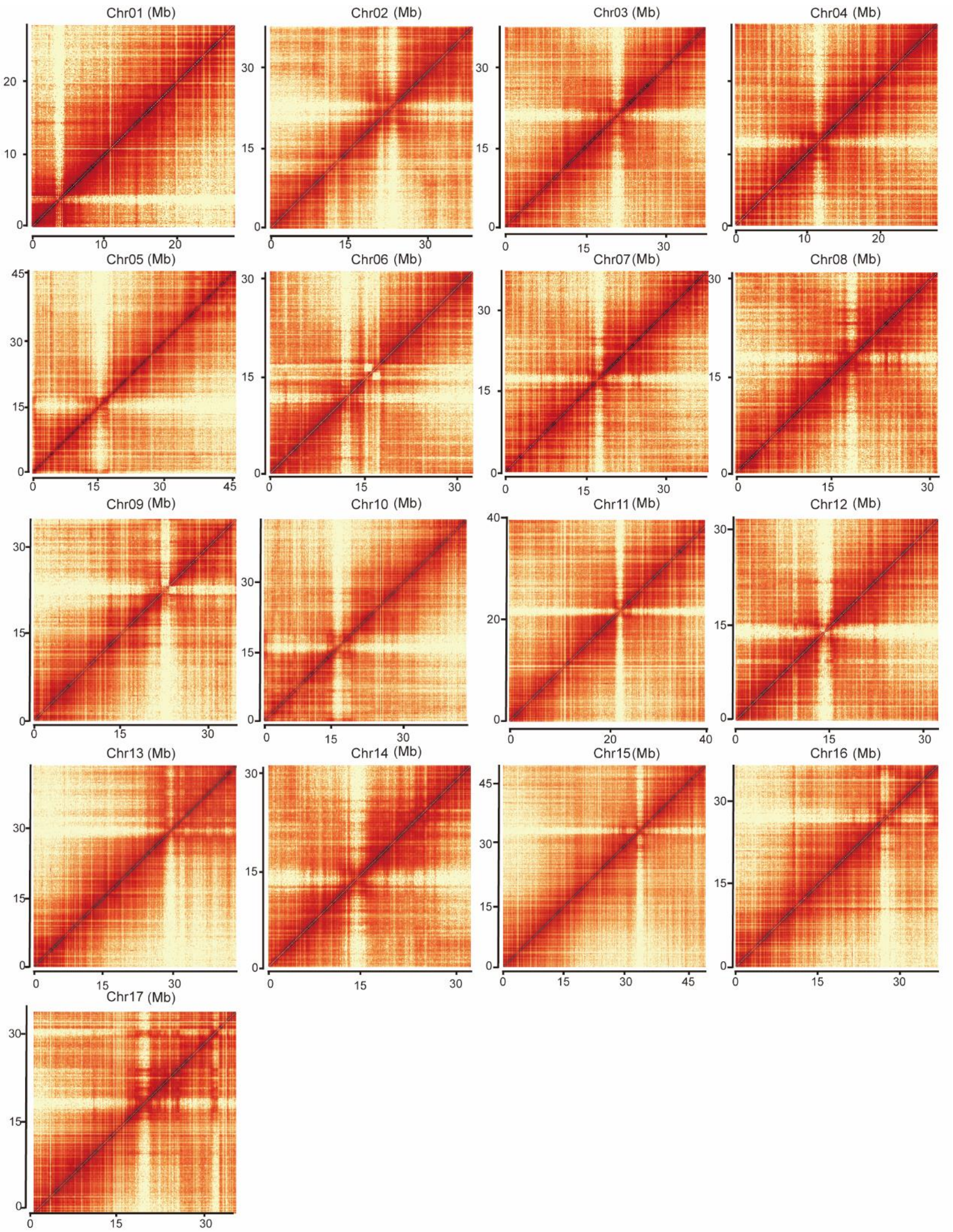


**Figure S2. Hi-C interaction heat map**

The 17 chromosome Hi-c heat map

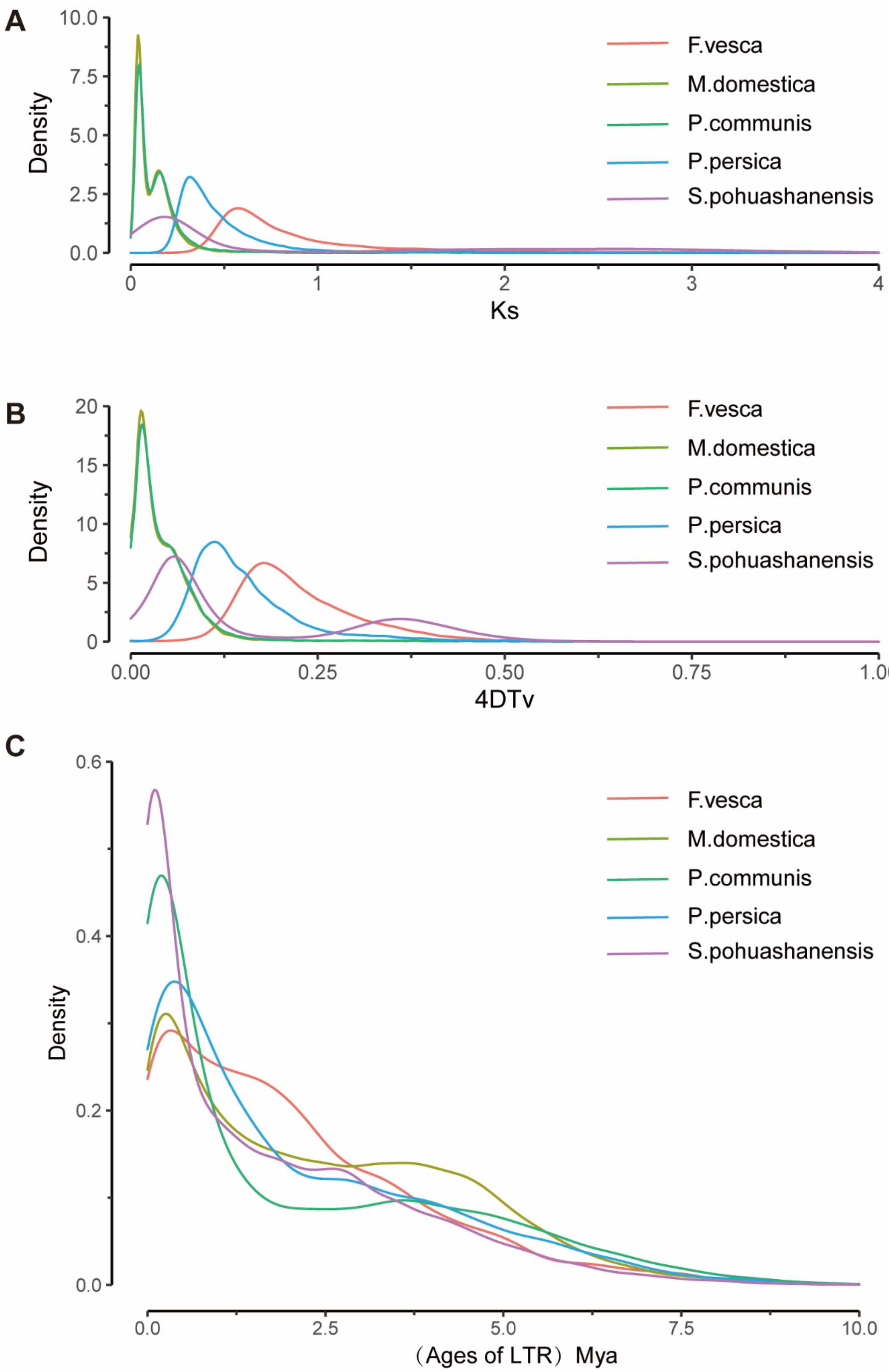


**Figure S3. Distribution of Ks, 4DTv and ages of LTR of *S. pohuashanensis* and other species**

a. Ks distribution of *S. pohuashanensis* and other representative species; b. 4DTv distribution of *S. pohuashanensis* and other representative species; c. Ages of LTR of *S. pohuashanensis* and other species (Molecular clock r is $7*10^{-9}$）.

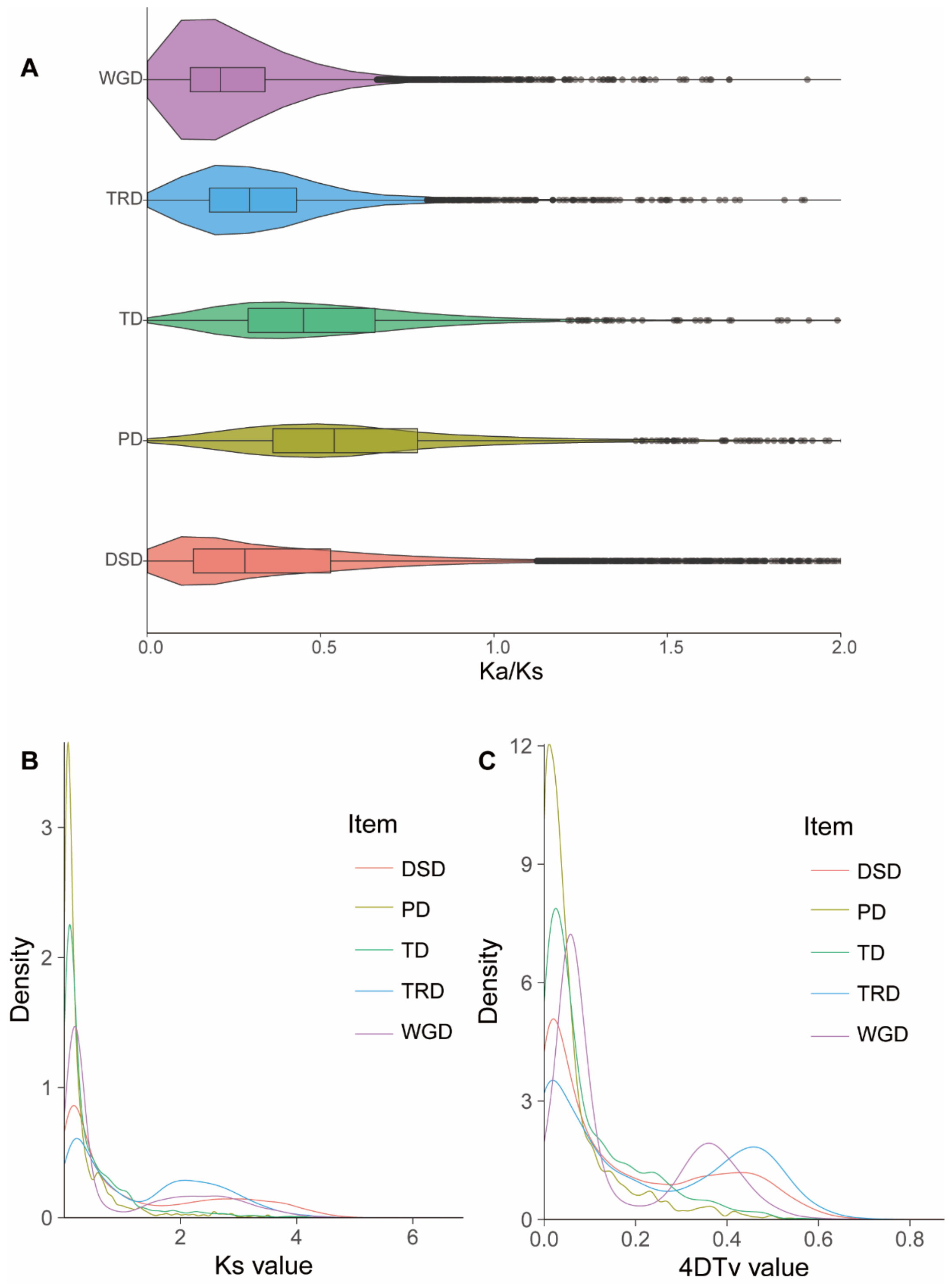


**Figure S4. Gene Duplication and evolution.**

**a**. Distribution of Ka/Ks of 5 replication types; **b**. Distribution of Ks of 5 replication types; **c**. Distribution of 4DTv of 5 replication types.

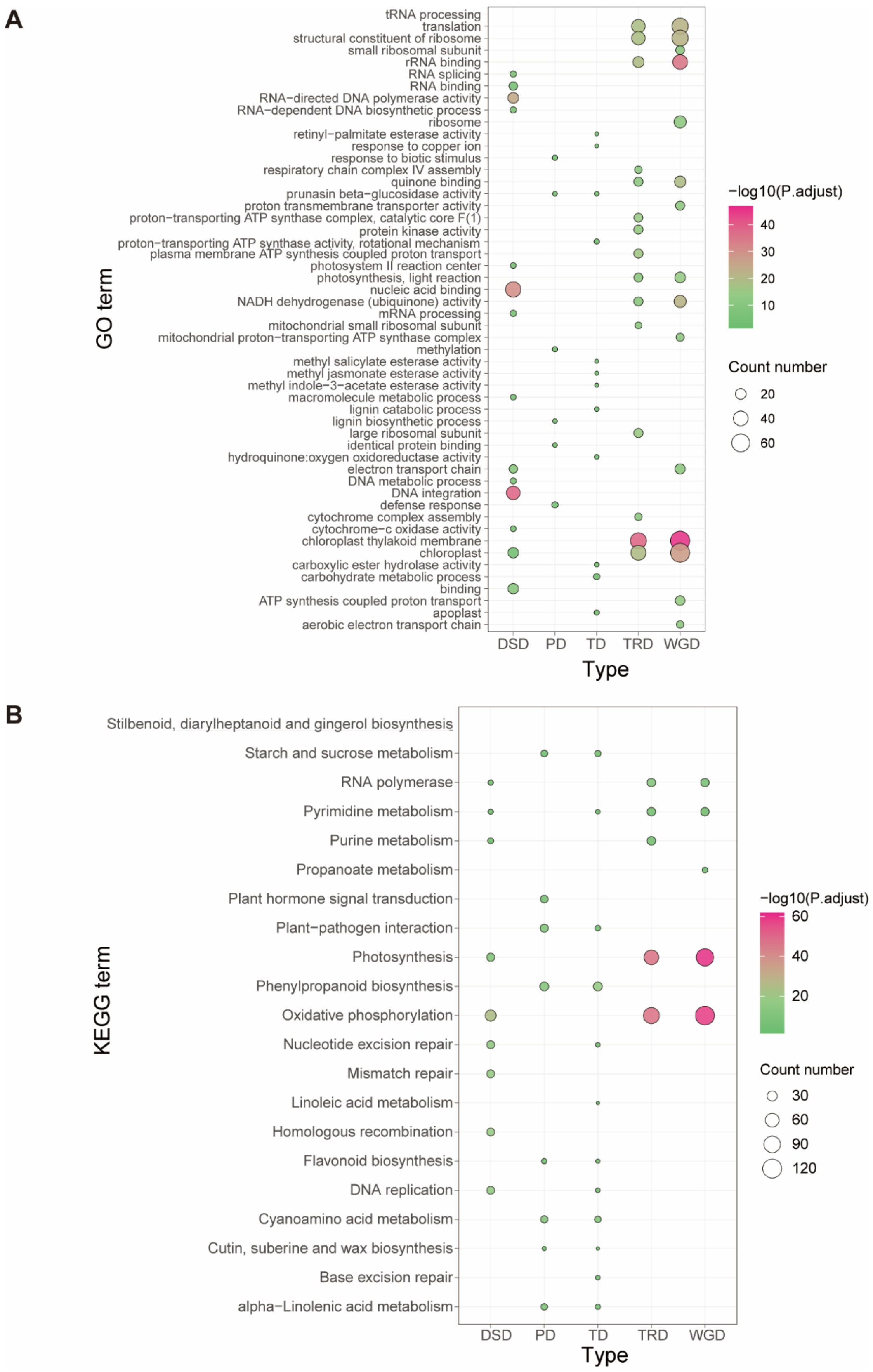


**Figure S5. Gene duplication enrichment analysis**

**a**. GO Enrichment Analysis of Genes Expanded in 5 Types of Replication; **b**. KEGG Enrichment Analysis of Genes Expanded in 5 Types of Replication

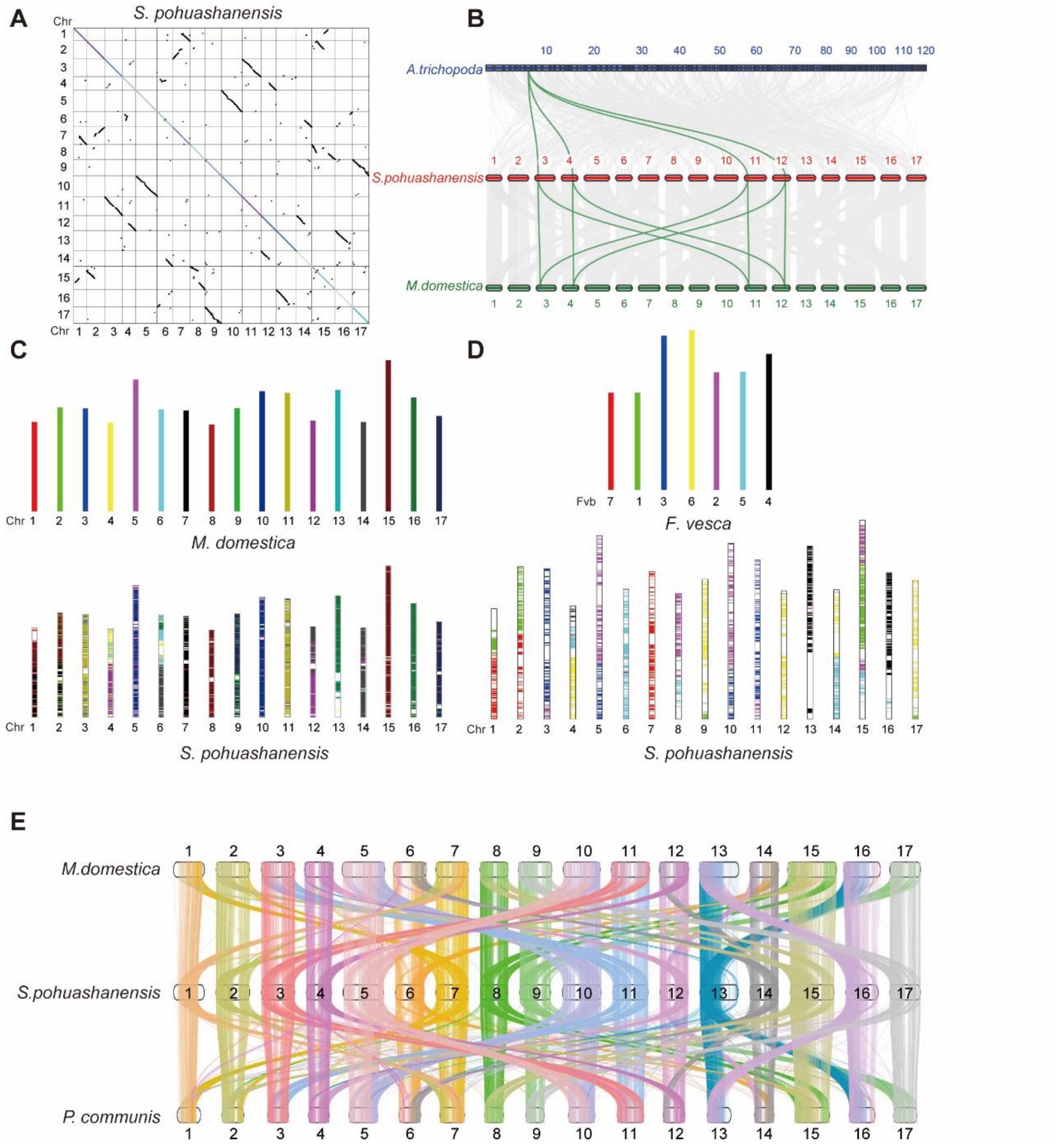


### Figure S6. Genome collinearity analysis

**a**. Dot plots of paralogs in the *S. pohuashanensis* genome; **b**. *A. trichopoda*, *S. pohuashanensis* and *M. domestica* gene level collinearity analysis; **c**. *S. pohuashanensis* and *M. domestica* gene level collinearity analysis; **d**. *Fragaria vesca and S. pohuashanensis* gene level collinearity analysis; **e**. *S. pohuashanensis, M. domestica* and *P. communis* genome level collinearity analysis;

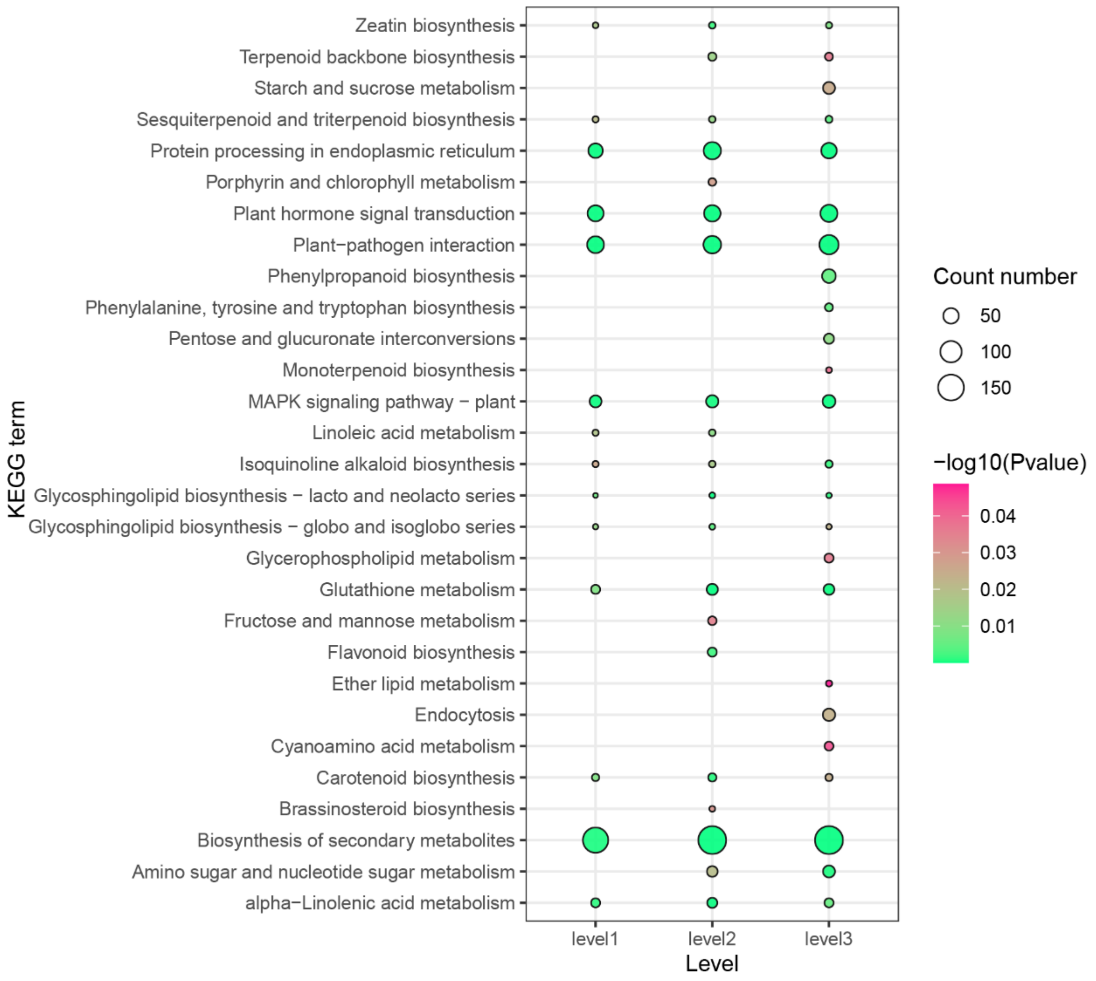


**Figure S7. KEGG enrichment pathways of 3 level DEGS**

The sunburn signaling pathways both enriched in the three stages are: biosynthesis of secondary metabolites, MAPK signaling pathway-plant, plant hormone signal transduction, plant-pathogen interaction.

**Supplementary Tables**

#### Table S1. Genome assembly statistic information for *S. pohuashanensis*

| | Type | Size(bp) | Number |
|---|---|---|---|
| scaffold | N10 | 45635594 | 2 |
| | N20 | 43700000 | 3 |
| | N30 | 39651545 | 5 |
| | N40 | 37451735 | 7 |
| | N50 | 36685546 | 8 |
| | N60 | 35059391 | 10 |
| | N70 | 32349783 | 12 |
| | N80 | 31915921 | 14 |
| | N90 | 28204096 | 16 |
| | smallest | 14040 | |
| | longest | 49422898 | |
| | total | 660101906 | 404 |
| contig | N10 | 43197690 | 2 |
| | N20 | 37451735 | 4 |
| | N30 | 36685546 | 5 |
| | N40 | 32349783 | 7 |
| | N50 | 28204096 | 10 |
| | N60 | 26182626 | 12 |
| | N70 | 20502803 | 15 |
| | N80 | 13843604 | 19 |
| | N90 | 11623214 | 24 |
| | smallest | 14040 | |
| | longest | 43700000 | |
| | total | 660099706 | 426 |
| Illumina | Total_reads | 267,610,056 | |
| | Mapped_reads | 263,855,425 | |
| | Mapped(%) | 98.6 | |
| | Properly_mapped_reads | 248,191,932 | |
| | Properly_mapped | 93.72% | |
| PacBio | Reads_num | 1,307,792 | |
| | Reads_base | 26.96Gb | |
| | Reads_LenN50 | 20.26Kb | |
| | Reads_LenMean | 20.62Kb | |
| | Reads_LenMax | 50.23Kb | |
| | Contig number | 414 | |
| | Contig length | 660.10Mb | |
| | Contig N50 | 32.35Mb | |
| | Contig N90 | 11.71Mb | |
| | Contig max | 44.08Mb | |
| | GC content | 38.22% | |

**Table S2. BUSCO evaluation result**

| Type | Number |
|---|---|
| Complete BUSCOs(C) | 2065 (97.36%) |
| Complete and single-copy BUSCOs(S) | 1178 (55.54%) |
| Complete and duplicated BUSCOs(D) | 887 (41.82%) |
| Fragmented BUSCOs(F) | 12 (0.57%) |
| Missing BUSCOs(M) | 44 (2.07%) |
| Total Lineage BUSCOs | 2121 |

**Table S3. Repeat sequence**

| Type | Number | Length | Rate(%) |
|---|---|---|---|
| ClassI | 762,895 | 358,607,246 | 54.33 |
| ClassI/DIRS | 21,823 | 15,317,593 | 2.32 |
| ClassI/LARD | 182,335 | 48,516,579 | 7.35 |
| ClassI/LINE | 55,413 | 17,638,794 | 2.67 |
| ClassI/LTR/Copia | 216,443 | 143,037,376 | 21.67 |
| ClassI/LTR/Gypsy | 221,788 | 146,523,609 | 22.2 |
| ClassI/LTR/Unknown | 57,739 | 19,611,637 | 2.97 |
| ClassI/PLE | 2,156 | 762,406 | 0.12 |
| ClassI/SINE | 2,205 | 426,058 | 0.06 |
| ClassI/TRIM | 2,193 | 1,348,654 | 0.2 |
| ClassI/Unknown | 800 | 354,697 | 0.05 |
| ClassII | 349,151 | 97,968,107 | 14.84 |
| ClassII/Crypton | 8 | 427 | 0 |
| ClassII/Helitron | 47,215 | 12,712,700 | 1.93 |
| ClassII/MITE | 1,990 | 354,701 | 0.05 |
| ClassII/Maverick | 812 | 2,460,120 | 0.37 |
| ClassII/TIR | 250,344 | 74,228,567 | 11.25 |
| ClassII/Unknown | 48,782 | 11,140,826 | 1.69 |
| PotentialHostGene | 8,998 | 9,605,880 | 1.46 |
| SSR | 547 | 86,788 | 0.01 |
| Unknown | 47,437 | 16,955,553 | 2.57 |
| Total | 1,169,028 | 451,799,456 | 68.44 |

**Table S4. Duplication gene's distribution**

| Types | NO. of gene pairs | NO. of genes | overlap with expand genes | expand genes |
|---|---|---|---|---|
| whole-genome duplication (WGD) | 18981 | 25604 (58.7%) | 912 | 4212 |
| tandem duplication (TD) | 1217 | 2903 (6.7%) | 439 | |
| proximal duplication (PD) | 1537 | 3462 (7.9%) | 670 | |

| | | | |
|---|---|---|---|
| transposed duplication (TRD) | 6593 | 5778 (13.2%) | 517 |
| dispersed duplication (DSD) | 13513 | 5886 (13.5%) | 1562 |

#### Table S5. Resequencing location (Summary of all accessions sequenced in this study)

| Accession ID | Origin | Longtiude (+E/-W) | Latitude (N) | ReadSum | Mapped ReadSum | Map Rate (%) | GC (%) | Q20 (%) | Q30 (%) |
|---|---|---|---|---|---|---|---|---|---|
| R01 | Mount Baihua, China | 117°57′19.624″ | 35°34′02.640″ | 54,595,302 | 53,160,164 | 97.37 | 37.99 | 96.83 | 91.5 |
| R02 | Moscow City, Russia | 37°40′56.190″ | 55°47′59.032″ | 59,191,224 | 56,456,398 | 95.38 | 37.86 | 97.52 | 93.14 |
| R03 | Mount, Kunyu, China | 121°44′39.631″ | 37°17′10.338″ | 58,064,702 | 52,203,035 | 89.9 | 37.73 | 96.84 | 91.58 |
| R04 | Mount Fanjing, China | 108°42′02.048″ | 27°55′10.754″ | 56,548,843 | 51,022,420 | 90.23 | 37.87 | 96.57 | 91.1 |
| R05 | Mount Tai, China | 117°6′08.712″ | 36°15′57.042″ | 63,021,322 | 61,779,466 | 98.03 | 37.76 | 97.57 | 93.23 |
| R06 | Mount Lao, China | 120°38′39.610″ | 36°12′27.630″ | 59,728,533 | 58,232,656 | 97.5 | 37.75 | 97.63 | 93.37 |
| R07 | Pangquan Ditch, China | 111°29′39.152″ | 37°50′50.082″ | 57,801,944 | 56,152,639 | 97.15 | 37.68 | 96.42 | 90.69 |
| R08 | Mount Tuoliang, China | 113°49′27.026″ | 38°43′40.508″ | 53,764,804 | 51,817,531 | 96.38 | 38.2 | 96.94 | 91.72 |
| R09 | Mount Baishi, China | 114°43′10.906″ | 39°13′52.658″ | 56,442,624 | 54,403,697 | 96.39 | 38.22 | 97.01 | 91.89 |
| R10 | Mount Wuling, China | 117°28′34.014″ | 40°38′01.378″ | 55,785,690 | 54,911,677 | 98.43 | 37.95 | 97.6 | 93.35 |
| R11 | Saihan Ba Forest Farm, China | 117°23′50.420″ | 42°22′00.404″ | 55,292,844 | 53,055,394 | 95.95 | 38.15 | 96.84 | 91.6 |
| R12 | Wangyedian Forest Farm, China | 118°48′08.957″ | 41°52′34.817″ | 58,625,766 | 57,617,259 | 98.28 | 37.82 | 97.68 | 93.5 |
| R13 | Haerbin City, China | 126°34′36.052″ | 45°44′57.422″ | 66,304,370 | 64,769,939 | 97.69 | 37.95 | 97.32 | 92.67 |
| R14 | Mount Ta, China | 118°6′00.000″ | 35°16′48.000″ | 59,425,663 | 58,173,052 | 97.89 | 37.76 | 97.57 | 93.2 |
| R15 | Wufeng County, China | 111°2′56.864″ | 30°11′14.154″ | 57,652,218 | 55,204,848 | 95.75 | 37.4 | 97.55 | 93.17 |

| | | | | | | | | | |
|---|---|---|---|---|---|---|---|---|---|
| R16 | Jinan City, China | 117°7′44.350″ | 36°40′36.545″ | 49,704,013 | 48,153,660 | 96.88 | 38.92 | 98.03 | 94.27 |
| R17 | Kew Royal Botanic Gardens, England | -0°17′36.560″ | 51°28′51.488″ | 47,015,608 | 46,019,597 | 97.88 | 38.85 | 97.53 | 92.94 |
| R18 | Mount Fanjing, China | 108°42′02.048″ | 27°55′10.754″ | 51,709,422 | 46,284,987 | 89.51 | 39.05 | 97.02 | 92.38 |
| R19 | Royal Botanic Gardens, Canada | -79°52′34.036″ | 43°17′23.719″ | 51,501,400 | 47,993,994 | 93.19 | 38.97 | 97.24 | 92.87 |
| R20 | Mount Fanjing, China | 108°42′02.048″ | 27°55′10.754″ | 49,864,285 | 45,531,963 | 91.31 | 38.19 | 98.28 | 94.83 |
| R21 | Mount, Kunyu, China | 121°44′39.631″ | 37°17′10.338″ | 50,659,018 | 47,279,497 | 93.33 | 38.51 | 96.71 | 91.87 |
| R22 | Mount Guangtu, China | 118°31′40.922″ | 41°19′34.752″ | 48,656,661 | 46,615,505 | 95.8 | 38.77 | 96.75 | 91.38 |

#### Table S6. Transcriptome sampling environmental data

| Type | Control | Level1 | Level2 | Level3 |
|---|---|---|---|---|
| Processing time (h) | 0 | 2.5 | 26.5 | 146.5 |
| Temperature (℃) | 35.08 | 40.78 | 39.05 | 39.33 |
| Brightness （Lx*1000） | 23.62 | 77.75 | 83.75 | 86.28 |
| Humidity （%） | 22.95 | 18.38 | 18.63 | 27.68 |

#### Table S7. Summary of categorized SNPs

| SNP category | No. SNPs |
|---|---|
| Upstream | 772,657 |
| Downstream | 641,663 |
| Upstream/downstream | 83,639 |
| Stop gain | 6,907 |
| Stop loss | 926 |
| Synonymous | 303,424 |
| Nonsynonymous | 378,589 |
| Intronic | 1,502,903 |
| Intergenic | 4,854,602 |
| UTR3 | 142,213 |
| UTR5 | 82,520 |
| Total | 8,770,043 |

#### Table S8. Annotation of 182 candidate genes in the response to sunburn

| Gene | Gene model | Description | $Log_2$ Fold change | Group | Up/Down |
|---|---|---|---|---|---|

| | | | | | |
|---|---|---|---|---|---|
| HSF | Sp12G001740.1 | heat shock transcription factor A3 | 1.18 | CK_L1 | Up |
| | Sp14G001710.1 | heat shock transcription factor A3 | 1.71 | CK_L1 | Up |
| | Sp04G001220.1 | heat shock transcription factor A5 | 1.12/1.16/1.18 | CK_L1/CK_L2/CK_L3 | Up |
| | Sp03G023500.1 | heat shock transcription factor A6b | 2.28/2.49 | CK_L2/CK_L3 | Up |
| | Sp11G025460.1 | heat shock transcription factor A6b | 2.58/2.42 | CK_L2/CK_L3 | Up |
| | Sp01G017660.1 | heat shock transcription factor B2a | 1.69/2.69/2.15 | CK_L1/CK_L2/CK_L3 | Up |
| | Sp07G023660.1 | heat shock transcription factor B2a | 1.88/1.09 | CK_L2/CK_L3 | Up |
| HSP20 | Sp01G020080.1 | HSP20 family protein | 6.00/6.01 | CK_L1/CK_L2 | Up |
| | Sp15G038330.1 | HSP20 family protein | 6.86/6.52 | CK_L1/CK_L2 | Up |
| | Sp09G019280.1 | HSP20 family protein | 5.13 | CK_L2 | Up |
| | Sp04G012430.1 | HSP20 family protein | 1.2 | CK_L3 | Up |
| | Sp17G024510.1 | HSP20 family protein | 6.11 | CK_L3 | Up |
| | Sp01G012780.1 | HSP20 family protein | 4.60/3.89/4.11 | CK_L1/CK_L2/CK_L3 | Up |
| | Sp05G022540.1 | HSP20 family protein | 7.39/6.18/6.28 | CK_L1/CK_L2/CK_L3 | Up |
| | Sp06G005610.1 | HSP20 family protein | 3.52/4.26/4.19 | CK_L1/CK_L2/CK_L3 | Up |
| | Sp07G019210.1 | HSP20 family protein | 7.01/4.89/3.83 | CK_L1/CK_L2/CK_L3 | Up |
| | Sp07G022390.1 | HSP20 family protein | 2.19/4.09/4.26 | CK_L1/CK_L2/CK_L3 | Up |
| | Sp07G026110.1 | HSP20 family protein | 4.41/4.53/3.60 | CK_L1/CK_L2/CK_L3 | Up |

| | | | | | |
|---|---|---|---|---|---|
| | Sp08G006210.1 | HSP20 family protein | 4.85/2.99/3.45 | CK_L1/CK_L2/CK_L3 | Up |
| | Sp08G006240.1 | HSP20 family protein | 10.97/9.15/6.47 | CK_L1/CK_L2/CK_L3 | Up |
| | Sp08G022710.1 | HSP20 family protein | 8.03/5.88/5.32 | CK_L1/CK_L2/CK_L3 | Up |
| | Sp10G016790.1 | HSP20 family protein | 11.84/11.04/7.52 | CK_L1/CK_L2/CK_L3 | Up |
| | Sp13G008820.1 | HSP20 family protein | 5.12/4.91/5.43 | CK_L1/CK_L2/CK_L3 | Up |
| | Sp14G008870.1 | HSP20 family protein | 8.38/6.46/8.27 | CK_L1/CK_L2/CK_L3 | Up |
| | Sp17G001570.1 | HSP20 family protein | 8.07/8.79/3.89 | CK_L1/CK_L2/CK_L3 | Up |
| | Sp17G013620.1 | HSP20 family protein | 3.64/4.42/4.15 | CK_L1/CK_L2/CK_L3 | Up |
| | Sp17G019110.1 | HSP20 family protein | 6.60/8.70/7.76 | CK_L1/CK_L2/CK_L3 | Up |
| HSP60 | Sp05G017640.1 | Heat shock protein 60 | 1.20/1.70/1.49 | CK_L1/CK_L2/CK_L3 | Up |
| HSP70 | Sp05G005390.1 | heat shock 70kDa protein 1/8 | 1.61/1.51 | CK_L2/CK_L3 | Up |
| | Sp01G011620.1 | heat shock 70kDa protein 1/8 | 2.34/2.78/2.66 | CK_L1/CK_L2/CK_L3 | Up |
| | Sp01G011670.1 | heat shock 70kDa protein 1/8 | 1.49/1.54/1.48 | CK_L1/CK_L2/CK_L3 | Up |
| | Sp02G000690.1 | heat shock 70kDa protein 1/8 | 5.05/4.54/5.18 | CK_L1/CK_L2/CK_L3 | Up |
| | Sp07G018300.1 | heat shock 70kDa protein 1/8 | 1.57/2.31/2.13 | CK_L1/CK_L2/CK_L3 | Up |
| | Sp07G018310.1 | heat shock 70kDa protein 1/8 | 1.17/1.92/1.43 | CK_L1/CK_L2/CK_L3 | Up |
| | Sp08G021980.1 | heat shock 70kDa protein 1/8 | 1.09/1.28/1.16 | CK_L1/CK_L2/CK_L3 | Up |
| | Sp12G019950.1 | heat shock 70kDa protein 5 | 4.23/3.64/2.39 | CK_L1/CK_L2/CK_L3 | Up |

| | | | | | |
|---|---|---|---|---|---|
| | Sp13G015930.1 | heat shock 70kDa protein 4 | 1.25/1.54/1.73 | CK_L1/CK_L2/CK_L3 | Up |
| | Sp15G012200.1 | heat shock 70kDa protein 1/8 | 2.82/2.46/2.08 | CK_L1/CK_L2/CK_L3 | Up |
| | Sp15G037680.1 | heat shock 70kDa protein 1/8 | 1.77/1.17/1.61 | CK_L1/CK_L2/CK_L3 | Up |
| | Sp16G016060.1 | heat shock 70kDa protein 4 | 2.12/2.60/1.52 | CK_L1/CK_L2/CK_L3 | Up |
| | Sp17G020560.1 | heat shock 70kDa protein 1/8 | 7.30/5.25/5.06 | CK_L1/CK_L2/CK_L3 | Up |
| | Sp17G020570.1 | heat shock 70kDa protein 1/8 | 4.92/3.91/4.90 | CK_L1/CK_L2/CK_L3 | Up |
| HSP90 | Sp15G000820.1 | heat shock protein 90kDa beta | 1.37 | CK_L2 | Up |
| | Sp11G003000.1 | heat shock protein 90-2 | 1.35/1.26 | CK_L2/CK_L3 | Up |
| | Sp13G004590.1 | heat shock protein 90kDa beta | 1.58/1.65 | CK_L2/CK_L3 | Up |
| | Sp17G010330.1 | heat shock protein 90kDa beta | 1.44/1.40 | CK_L2/CK_L3 | Up |
| DREB | Sp04G014700.1 | Dehydration-responsive element-binding protein (DREB2) | 2.62/2.05 | CK_L1/CK_L2 | Up |
| | Sp12G016790.1 | Dehydration-responsive element-binding protein (DREB2) | 1.95/2.44/1.70 | CK_L1/CK_L2/CK_L3 | Up |
| | Sp07G023450.1 | DREB protein | -0.199225685 | CK_L1/CK_L2/CK_L3 | Down |
| | Sp15G018170.1 | Dehydration-responsive element-binding protein (DREB5) | 1.261682243 | CK_L1/CK_L3 | Down |

| | | | | | |
|---|---|---|---|---|---|
| | Sp01G008730.1 | NAC domain-containing protein 72-like | 5.25 | CK_L1 | Up |
| | Sp01G008750.1 | NAC domain-containing protein 102-like | 1.9 | CK_L1 | Up |
| | Sp02G020780.1 | NAC domain class transcription factor | 1.02 | CK_L1 | Up |
| | Sp07G015450.1 | NAC domain-containing protein 68-like | 1.94 | CK_L1 | Up |
| | Sp14G014970.1 | NAC transcription factor 29-like | 1.46 | CK_L1 | Up |
| | Sp01G000310.1 | NAC domain protein | 1.08 | CK_L2 | Up |
| | Sp06G013180.1 | NAC transcription factor 25-like | 1.07 | CK_L2 | Up |
| NAC | Sp11G014660.1 | NAC domain-containing protein 90-like | 1.32 | CK_L2 | Up |
| | Sp04G003360.1 | NAC domain-containing protein 2-like | 1.96/1.73 | CK_L1/CK_L2 | Up |
| | Sp07G015480.1 | NAC domain-containing protein 72-like | 2.16/1.08 | CK_L1/CK_L2 | Up |
| | Sp07G006120.1 | NAC domain class transcription factor | 1.29/1.01 | CK_L1/CK_L3 | Up |
| | Sp10G017840.1 | NAC domain protein | 1.87/2.23 | CK_L2/CK_L3 | Up |
| | Sp01G008860.1 | putative NAC domain-containing protein 94 | 3.14/3.16/1.89 | CK_L1/CK_L2/CK_L3 | Up |
| | Sp01G008770.1 | NAC domain-containing protein 102-like | 1.42/1.67/1.19 | CK_L1/CK_L2/CK_L3 | Up |

| | | | | | |
|---|---|---|---|---|---|
| | Sp01G008680.1 | NAC domain-containing protein 74-like | 2.22 | CK_L1 | Down |
| | Sp03G013180.1 | NAC domain-containing protein 90 isoform X2 | -1.79 | CK_L1 | Down |
| | Sp11G014820.1 | NAC domain-containing protein 89-like | -1.74 | CK_L1 | Down |
| | Sp15G011170.1 | NAC domain protein | -1.06 | CK_L1 | Down |
| | Sp12G003010.1 | NAC domain-containing protein 90-like | 0.841549296 | CK_L1/CK_L3 | Down |
| | Sp16G005830.1 | NAC domain class transcription factor | 1.342342342 | CK_L1/CK_L2 | Down |
| | Sp01G020310.1 | NAC domain class transcription factor | 0.862318841 | CK_L2/CK_L3 | Down |
| | Sp05G013710.1 | NAC transcription factor 29-like | -0.632852919 | CK_L1/CK_L2/CK_L3 | Down |
| | Sp07G026570.1 | NAC domain class transcription factor | -0.309249794 | CK_L1/CK_L2/CK_L3 | Down |
| | Sp09G001730.1 | NAC domain class transcription factor | -1.070234114 | CK_L1/CK_L2/CK_L3 | Down |
| | Sp10G013130.1 | NAC transcription factor 29-like | -0.63328618 | CK_L1/CK_L2/CK_L3 | Down |
| | Sp17G000460.1 | NAC domain class transcription factor | -0.648422091 | CK_L1/CK_L2/CK_L3 | Down |
| MYC | Sp06G011680.1 | transcription factor MYC3-like | 1.190217391 | CK_L1/CK_L3 | Down |

| | | | | | |
|---|---|---|---|---|---|
| | Sp16G02385 0.1 | transcription factor MYC2 | 0.919075145 | CK_L1/CK_L3 | Down |
| | Sp01G00809 0.1 | transcription factor MYC2 | -0.250276167 | CK_L1/CK_L2/CK _L3 | Down |
| | Sp01G00808 0.1 | transcription factor MYC2-like | -0.427731242 | CK_L1/CK_L2/CK _L3 | Down |
| | Sp06G01167 0.1 | transcription factor MYC2-like | -0.7951988 | CK_L1/CK_L2/CK _L3 | Down |
| | Sp06G00292 0.1 | transcription factor MYC2 | -0.445036916 | CK_L1/CK_L2/CK _L3 | Down |
| | Sp14G01348 0.1 | transcription factor MYC3-like isoform X2 | -0.319952444 | CK_L1/CK_L2/CK _L3 | Down |
| | Sp14G01352 0.1 | transcription factor MYC2-like | -0.121865661 | CK_L1/CK_L2/CK _L3 | Down |
| MYB | Sp15G01727 0.1 | myb proto-oncogene protein | 1.25/1.14/1.1 8 | CK_L1/CK_L2/CK _L3 | Up |
| | Sp11G02928 0.1 | myb proto-oncogene protein | 1.28/2.18/1.7 4 | CK_L1/CK_L2/CK _L3 | Up |
| | Sp02G00650 0.1 | myb proto-oncogene protein | 1.06/1.03/1.2 0 | CK_L1/CK_L2/CK _L3 | Up |
| | Sp03G02689 0.1 | myb proto-oncogene protein | 1.55/2.13/1.7 6 | CK_L1/CK_L2/CK _L3 | Up |
| | Sp01G00180 0.1 | transcription factor MYB90-like isoform X1 | 2.56/3.88/2.7 9 | CK_L1/CK_L2/CK _L3 | Up |
| | Sp11G00969 0.1 | myb proto-oncogene protein | 1.13/1.77/1.8 1 | CK_L1/CK_L2/CK _L3 | Up |
| | Sp07G01464 0.1 | myb proto-oncogene protein | 1.28/2.88/1.6 1 | CK_L1/CK_L2/CK _L3 | Up |
| | Sp16G00315 0.1 | myb proto-oncogene protein | 1.08/1.56/1.1 1 | CK_L1/CK_L2/CK _L3 | Up |

| | | | | | |
|---|---|---|---|---|---|
| | Sp14G01780 0.1 | myb proto-oncogene protein | 1.32/2.75/1.72 | CK_L1/CK_L2/CK_L3 | Up |
| | Sp08G00686 0.1 | myb proto-oncogene protein | 1.95/1.93 | CK_L1/CK_L3 | Up |
| | Sp07G02261 0.1 | myb proto-oncogene protein | 1.55/1.62 | CK_L1/CK_L3 | Up |
| | Sp08G00816 0.1 | myb proto-oncogene protein | -0.319352778 | CK_L1/CK_L2/CK_L3 | Down |
| | Sp15G00625 0.1 | myb proto-oncogene protein | -0.730593607 | CK_L1/CK_L2/CK_L3 | Down |
| | Sp05G01019 0.1 | myb proto-oncogene protein | -0.239674955 | CK_L1/CK_L2/CK_L3 | Down |
| | Sp02G01471 0.1 | myb proto-oncogene protein | -0.271922977 | CK_L1/CK_L2/CK_L3 | Down |
| | Sp15G02395 0.1 | myb proto-oncogene protein | -0.840930852 | CK_L1/CK_L2/CK_L3 | Down |
| | Sp09G01575 0.1 | myb proto-oncogene protein | 0.69047619 | CK_L1/CK_L3 | Down |
| | Sp05G01920 0.1 | myb proto-oncogene protein | -0.43252726 | CK_L1/CK_L2/CK_L3 | Down |
| | Sp09G00817 0.1 | myb proto-oncogene protein | -0.509409371 | CK_L1/CK_L2/CK_L3 | Down |
| | Sp17G00800 0.1 | myb proto-oncogene protein | 1.190045249 | CK_L1/CK_L3 | Down |
| | Sp17G02491 0.1 | myb proto-oncogene protein | -0.732600733 | CK_L1/CK_L2/CK_L3 | Down |
| | Sp04G01636 0.1 | myb proto-oncogene protein | -0.723385187 | CK_L1/CK_L2/CK_L3 | Down |
| ERF | Sp07G02199 0.1 | ethylene-responsive | 1.345744681 | CK_L1/CK_L3 | Down |

| | | | | |
|---|---|---|---|---|
| | transcription factor ERF105-like | | | |
| Sp07G022000.1 | ethylene-responsive transcription factor ERF106-like | -0.494829805 | CK_L1/CK_L2/CK_L3 | Down |
| Sp01G017430.1 | ethylene-responsive transcription factor ERF027-like | -0.147218348 | CK_L1/CK_L2/CK_L3 | Down |
| Sp02G007480.1 | ethylene-responsive transcription factor ERF017 | 1.632716049 | CK_L1/CK_L3 | Down |
| Sp05G009400.1 | ethylene-responsive transcription factor ERF109-like | 1.334319527 | CK_L1/CK_L3 | Down |
| Sp06G010450.1 | ethylene-responsive transcription factor ERF061 | -0.644257703 | CK_L1/CK_L2/CK_L3 | Down |
| Sp01G015520.1 | ethylene-responsive transcription factor ERF105-like | -1.04 | CK_L1 | Down |
| Sp03G026470.1 | ethylene-responsive transcription factor ERF073-like | -0.637540773 | CK_L1/CK_L2/CK_L3 | Down |
| Sp15G002980.1 | ethylene-responsive transcription factor ERF003-like | -0.406264058 | CK_L1/CK_L2/CK_L3 | Down |
| Sp10G009240.1 | ethylene-responsive transcription | 0.718823529 | CK_L1/CK_L3 | Down |

| | | | | | |
|---|---|---|---|---|---|
| | | factor ERF109-like | | | |
| | Sp07G014490.1 | ethylene-responsive transcription factor ERF023-like | -0.447040162 | CK_L1/CK_L2/CK_L3 | Down |
| | Sp15G023320.1 | ethylene-responsive transcription factor ERF109-like | -0.042834898 | CK_L1/CK_L2/CK_L3 | Down |
| | Sp15G029920.1 | ethylene-responsive transcription factor ERF012-like | 0.685763889 | CK_L1/CK_L3 | Down |
| WRKY | Sp07G014010.1 | WRKY transcription factor 46 | -0.651692959 | CK_L1/CK_L2/CK_L3 | Down |
| | Sp01G007260.1 | Probable WRKY transcription factor 46 | 1.980392157 | CK_L1/CK_L2 | Down |
| | Sp07G021060.1 | probable WRKY transcription factor 70 | -1.119402985 | CK_L1/CK_L2/CK_L3 | Down |
| | Sp15G003170.1 | WRKY transcription factor 40b | -0.489252195 | CK_L1/CK_L2/CK_L3 | Down |
| | Sp15G004390.1 | probable WRKY transcription factor 50 | -0.711805556 | CK_L1/CK_L2/CK_L3 | Down |
| | Sp04G014830.1 | WRKY transcription factor 33 | -0.517742642 | CK_L1/CK_L2/CK_L3 | Down |
| | Sp04G015500.1 | probable WRKY transcription factor 38 | 1.768361582 | CK_L1/CK_L2 | Down |

| | | | | |
|---|---|---|---|---|
| Sp12G017000.1 | WRKY transcription factor 33 | -0.56176288 | CK_L1/CK_L2/CK_L3 | Down |
| Sp15G027970.1 | probable WRKY transcription factor 51 | -1.101214575 | CK_L1/CK_L2/CK_L3 | Down |
| Sp02G014600.1 | isoform X1 WRKY domain class transcription factor | -0.675675676 | CK_L1/CK_L2/CK_L3 | Down |
| Sp08G011130.1 | WRKY domain class transcription factor | 1.025423729 | CK_L1/CK_L2 | Down |
| Sp11G004870.1 | WRKY transcription factor 33 | -0.948717949 | CK_L1/CK_L2/CK_L3 | Down |
| Sp16G020880.1 | WRKY domain class transcription factor | -0.960543816 | CK_L1/CK_L2/CK_L3 | Down |
| Sp15G008600.1 | WRKY domain class transcription factor | 0.992592593 | CK_L1/CK_L2 | Down |
| Sp06G010330.1 | probable WRKY transcription factor 53 | -0.260058588 | CK_L1/CK_L2/CK_L3 | Down |
| Sp01G014830.1 | probable WRKY transcription factor 70 | -2.566911197 | CK_L1/CK_L2/CK_L3 | Down |
| Sp12G017750.1 | probable WRKY transcription factor 70 isoform X2 | -0.132989632 | CK_L1/CK_L2/CK_L3 | Down |
| Sp13G020400.1 | WRKY domain class transcription factor | 0.861538462 | CK_L1/CK_L3 | Down |

| | | | | | |
|---|---|---|---|---|---|
| | Sp08G005390.1 | WRKY transcription factor 40b | 0.971698113 | CK_L1/CK_L2 | Down |
| | Sp05G032810.1 | WRKY transcription factor 42 | 1.037383178 | CK_L1/CK_L2 | Down |
| | Sp06G017210.1 | probable WRKY transcription factor 61 | 1.040540541 | CK_L1/CK_L2 | Down |
| | Sp14G012190.1 | probable WRKY transcription factor 53 | 1.963414634 | CK_L1/CK_L2 | Down |
| | Sp13G012330.1 | WRKY domain class transcription factor | -1.24 | CK_L1 | Down |
| | Sp09G019940.1 | WRKY transcription factor 40a | -1.06 | CK_L1 | Down |
| | Sp15G023780.1 | probable WRKY transcription factor 15 | -1.01 | CK_L1 | Down |
| | Sp01G006540.1 | WRKY transcription factor 22 | 0.560846561 | CK_L1/CK_L3 | Down |
| TIFY | Sp09G015320.1 | protein TIFY 11B-like | -0.217407026 | CK_L1/CK_L2/CK_L3 | Down |
| | Sp13G001760.1 | protein TIFY 6B-like | -0.37444468 | CK_L1/CK_L2/CK_L3 | Down |
| | Sp02G007440.1 | protein TIFY 10A | -0.350063648 | CK_L1/CK_L2/CK_L3 | Down |
| | Sp10G024460.1 | protein TIFY 5A | -0.108958202 | CK_L1/CK_L2/CK_L3 | Down |
| | Sp13G010420.1 | protein TIFY 9 | -0.216246554 | CK_L1/CK_L2/CK_L3 | Down |
| | Sp17G014810.1 | protein TIFY 10B-like | -0.184398431 | CK_L1/CK_L2/CK_L3 | Down |
| | Sp05G026960.1 | protein TIFY 5A-like | -0.11982124 | CK_L1/CK_L2/CK_L3 | Down |
| | Sp16G010480.1 | protein TIFY 9-like | -0.392761261 | CK_L1/CK_L2/CK_L3 | Down |

| | | | | | |
|---|---|---|---|---|---|
| | Sp07G008190.1 | protein TIFY 6b-like | -0.280373832 | CK_L1/CK_L2/CK_L3 | Down |
| | Sp15G018130.1 | protein TIFY 10A-like | -0.34379314 | CK_L1/CK_L2/CK_L3 | Down |
| | Sp05G026970.1 | protein TIFY 5A-like | -0.145347013 | CK_L1/CK_L2/CK_L3 | Down |
| | Sp10G024450.1 | protein TIFY 5A-like | -0.118864081 | CK_L1/CK_L2/CK_L3 | Down |
| MBF 1c | Sp15G015520.1 | multiprotein-bridging factor 1c-like | 3.01/3.49/3.80 | CK_L1/CK_L2/CK_L3 | Up |
| | Sp02G004270.1 | multiprotein-bridging factor 1c-like | 1.87/2.82/2.86 | CK_L1/CK_L2/CK_L3 | Up |
| ELIPs | Sp13G016710.1 | early light-induced protein 1 | 4.41/5.65/4.18 | CK_L1/CK_L2/CK_L3 | Up |
| | Sp13G016760.1 | early light-induced protein 1 | 4.11/5.69/4.06 | CK_L1/CK_L2/CK_L3 | Up |
| | Sp14G015020.1 | early light-induced protein 1 | 5.73/7.29/5.99 | CK_L1/CK_L2/CK_L3 | Up |
| | Sp13G016690.1 | early light-induced protein 1 | 7.46 | CK_L2 | Up |
| | Sp13G016740.1 | early light-induced protein 1 | 9.17 | CK_L2 | Up |
| | Sp13G016650.1 | early light-induced protein 1 | 5.5 | CK_L2 | Up |
| PAL | Sp12G011060.1 | phenylalanine ammonia-lyase 1 | 1.04/1.47 | CK_L1/CK_L2 | Up |
| | Sp04G009140.1 | phenylalanine ammonia-lyase 1-like | 1.08/1.49/1.03 | CK_L1/CK_L2/CK_L3 | Up |
| | Sp01G009770.1 | phenylalanine ammonia-lyase 1 | -1.18 | CK_L3 | Down |
| 4CL | Sp06G014010.1 | 4-coumarate--CoA ligase-like 9 | 1.87/1.99/2.14 | CK_L1/CK_L2/CK_L3 | Up |

| | | | | | |
|---|---|---|---|---|---|
| | Sp03G010890.1 | 4-coumarate--CoA ligase | -1.28 | CK_L1 | Down |
| | Sp03G010880.1 | 4-coumarate--CoA ligase | -1.64 | CK_L2 | Down |
| | Sp11G012280.1 | 4-coumarate--CoA ligase | -1.01 | CK_L2 | Down |
| | Sp16G009360.1 | 4-coumarate--CoA ligase-like 9 | -1.21 | CK_L2 | Down |
| | Sp13G021690.1 | 4-coumarate--CoA ligase | -0.567923671 | CK_L1/CK_L2/CK_L3 | Down |
| | Sp16G009350.1 | 4-coumarate--CoA ligase-like 9 | -0.61842919 | CK_L1/CK_L2/CK_L3 | Down |
| CHS | Sp04G000210.1 | chalcone synthase | 2.40/1.39 | CK_L2/CK_L3 | Up |
| | Sp04G000200.1 | chalcone synthase | 2.47/1.21 | CK_L2/CK_L3 | Up |
| | Sp04G000190.1 | chalcone synthase | 1.48 | CK_L2 | Up |
| FLS | Sp10G019790.1 | Molecular Function: flavonol synthase activity (GO:0045431) | 7.66/5.65 | CK_L1/CK_L2 | Up |
| | Sp01G013480.1 | flavonol synthase/flavanone 3-hydroxylase-like | -0.751373383 | CK_L1/CK_L2/CK_L3 | Down |
| | Sp06G012060.1 | flavonol synthase/flavanone 3-hydroxylase-like | 0.99380805 | CK_L1/CK_L3 | Down |
| | Sp11G013810.1 | flavonol synthase 3-like | -3.27 | CK_L1 | Down |
